# Graph-based pangenome provides insights into the adaptive evolution of *Cucurbita pepo*

**DOI:** 10.64898/2026.05.27.728270

**Authors:** Xuebo Zhao, Kailiang Bo, Honghe Sun, Jie Zhang, Arthur A. Schaffer, Adi Faigenboim, Harry S. Paris, Elad Oren, Amit Gur, Rebecca Grumet, Changlin Wang, Yong Xu, Zhangjun Fei

**Author notes:** These authors contributed equally (Xuebo Zhao, Kailiang Bo).

## Abstract

Understanding how crops adapt to environmental variation is crucial for biodiversity conservation and food security. *Cucurbita pepo* (pumpkin, squash, gourd), one of the earliest domesticated crops, exhibits remarkable phenotypic and ecological diversity, providing a valuable system for investigating the genomic basis of adaptation. Here, we constructed a graph-based *C. pepo* pangenome using nine chromosome-level assemblies and identified 229,431 high-confidence structural variants (SVs), which were genotyped across 206 wild and cultivated accessions. We reveal a history of parallel domestication, in which two deeply diverged gene pools independently gave rise to the cultivated *pepo* and *ovifera* lineages. Subsequent expansion into diverse environments was accompanied by pronounced genomic differentiation, with adaptive signatures largely associated with evolutionarily young alleles. Single-nucleotide polymorphisms (SNPs) and SVs contribute complementary dimensions of environmental responsiveness. Biogeographical modeling predicts further range contractions of wild relatives and elevated genetic offset in Northern North American populations under projected future climates. Notably, populations with higher genetic load harbor fewer climate-associated variants and exhibit greater predicted genetic offset, revealing an association between deleterious mutation burden and reduced adaptive potential. These findings highlight how genomic variation, adaptive diversity, and genetic load together shape environmental adaptation and inform conservation and crop improvement.

## Introduction

Environmental variation shapes species distributions, influences biodiversity, and affects global food security^1^. As environmental conditions deviate from historical norms, both natural populations and agricultural species face increasing challenges in maintaining productivity, persistence, and local adaptation. These pressures are particularly pronounced for crops, where rising global demand coincides with diverse environmental constraints, underscoring the urgency of developing well-adapted varieties^2^. Understanding how plants respond to varying environments and how evolutionary potential is distributed across cultivated crops and their wild relatives, is therefore critical for harnessing genetic resources to sustain agricultural productivity^3^.

Population genomic studies have provided valuable insights into environment-associated genetic variation^4,5^, yet most genomic analyses focus primarily on single-nucleotide polymorphisms (SNPs), which represent fine-scale variation but fail to capture large-effect structural variants (SVs) that can strongly influence gene function and phenotype^6^. SVs‒including insertions, deletions, inversions, and translocations‒constitute a major component of genomic diversity and play critical roles in phenotypic plasticity and environmental adaptation^7^. Recent advances in sequencing technologies and computational approaches have enabled large-scale SV identification, providing new opportunities to investigate their functional significance^8^. However, the contribution of SVs to local adaptation and their potential role in predicting the responses of species to future climates remain poorly understood. Moreover, adaptive potential is shaped not only by adaptive variants but also by genome-wide genetic load, raising the unresolved question of how deleterious mutations interact with adaptive diversity to constrain or facilitate environmental responses. Integrating SNPs and SVs within predictive genomic frameworks therefore provides a valuable opportunity to link genomic architecture, adaptive variation, and projected future environmental conditions^9^.

*Cucurbita pepo* L. is one of the first species to have been domesticated, with archaeological evidence suggestive of domestication dating back 10,000 years^10^. This species encompasses a wide range of cultivated, edible forms, including pumpkins and summer and winter squash, and is of considerable economic and nutritional importance^11,12^. Now cultivated globally across tropical and temperate regions, *C. pepo* exhibits remarkable morphological and genetic diversity^13^. Its long domestication history^14^ and broad ecological distribution spanning diverse environments^15^ make this crop species a powerful system for dissecting the genomic basis of environmental adaptation.

Here, we combined high-quality genome assemblies and population resequencing to construct a graph-based pangenome for *C. pepo*, enabling comprehensive identification and genotyping of SVs across diverse wild and cultivated accessions. By integrating genomic variants, including both SVs and SNPs, with environmental gradients and future climate projections, we investigate the genetic basis of local adaptation, assess population vulnerability under projected future climates, and evaluate how genetic load may constrain adaptive potential. Together, this work establishes a predictive framework that links genomic variation and adaptive potential, providing insights into the evolutionary dynamics of *C. pepo* and guiding germplasm conservation and crop improvement.

## Results

### Chromosome-scale genome assembly and gene-based pangenome construction

To enable accurate identification of SVs, we generated high-quality reference genomes for nine *C. pepo* accessions using PacBio HiFi long-read sequencing. This panel included the two major subspecies, *C. pepo* subsp. *ovifera* and subsp. *pepo*, comprising four accessions of *C. pepo* subsp. *ovifera* var. *ovifera* and five accessions of *C. pepo* subsp. *pepo* (**Fig. 1a** and **Supplementary Table 1**). In total, 97.03 Gb of HiFi data were produced, with an average of 10.78 Gb per accession (30.71× coverage). Using these data, we successfully assembled chromosome-scale genomes for all nine accessions. Final assembly sizes ranged from 332.1 Mb to 378.55 Mb (average 349.81 Mb), with contig N50 values between 3.51 Mb and 10.73 Mb (average 6.82 Mb) (**Supplementary Table 1**). On average, 93.85% of contigs were anchored to the 20 chromosomes, with unanchored sequences primarily composed of ribosomal DNA and other repetitive elements. BUSCO^16^ analysis revealed an average completeness of 98.06% of these assemblies, while k-mer based evaluation using Merqury^17^ showed an average completeness of 97.48% and quality values exceeding 60.74, corresponding to approximately one base error per million bases. The long terminal repeat (LTR) assembly index (LAI) ranged from 9.56 to 12.78 (average 10.98), meeting the criteria for reference-level assemblies^18^. Collectively, these metrics confirmed the high accuracy and completeness of the assemblies. Between 30,412 and 32,484 protein-coding genes were annotated per genome (average 31,501), with BUSCO completeness values ranging from 95.20% to 97.90% (average 96.94%) across the protein-coding gene sets (**Supplementary Table 1**). Repetitive sequences accounted for an average of 53.50% of the assemblies, ranging from 51.35% to 56.22% (**Supplementary Table 2**). Comparative genomic analysis revealed extensive macrosynteny among the nine assemblies, while large-scale chromosomal rearrangements including inversions and translocations were detected on chromosomes 6, 8, 11, and 13 (**Fig. 1a**).

**Figure 1.**
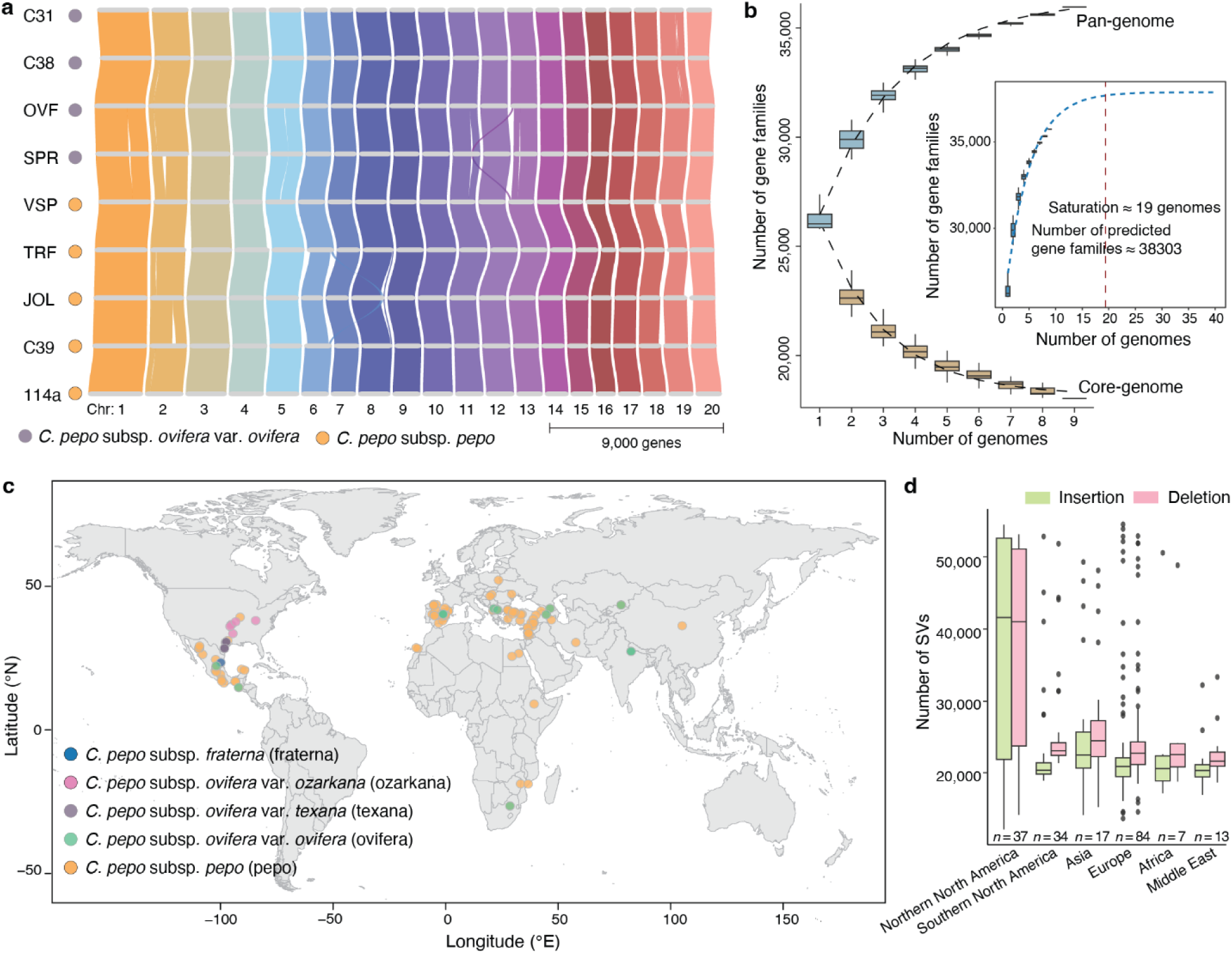
Pangenome of *C. pepo*. **a**, Chromosome-scale synteny map across the nine *C. pepo* assemblies, generated using GENESPACE (https://github.com/jtlovell/GENESPACE). **b**, Simulations of pangenome expansion and core-genome contraction. For each number of genomes, 100 random combinations were sampled from the nine genomes, and the numbers of non-redundant gene families in the pangenome and gene families shared across all sampled genomes in the core genome were calculated. Boxplots show the distributions across the 100 resampling iterations. The inset illustrates the predicted pangenome size and estimated saturation. **c**, Geographic distribution of 206 resequenced *C. pepo* accessions. Each colored dot represents an accession. The map was generated using ggplot2 (https://ggplot2.tidyverse.org/). **d,** Number of SVs identified in different *C. pepo* populations. For each boxplot, the lower and upper bounds indicate the first and third quartiles, respectively, the center line indicates the median, and the whiskers extend to 1.5× the interquartile range.

Using protein-coding genes from these nine assemblies, we constructed a gene-based pangenome to characterize gene presence–absence variation. Clustering of all protein-coding genes identified 35,956 distinct gene families, of which 18,041 (50.2%) were present in all nine genomes, 2,746 (7.6%) in eight genomes, 12,091 (33.6%) in two to seven genomes, and 3,078 (8.6%) in only a single genome. The accumulation of new gene families increased rapidly with the addition of the first few genomes and gradually approached a plateau. A nonlinear regression model fitted to the accumulation curve estimated that the pangenome would approach saturation at approximately 19 genomes, corresponding to around 38,303 gene families, indicating that our current sampling captured most of the diversity in *C. pepo* (∼93.87%) (**Fig. 1b**).

### Population-scale variants revealed by graph-based pangenome

Leveraging the advantages of graph-based pangenomes in capturing complex genomic variation^19^, we constructed a graph-based pangenome for *C. pepo* using nine chromosome-scale assemblies. This framework enabled the identification of 229,431 non-redundant SVs (≥20 bp), including 131,373 insertions and 98,058 deletions, collectively spanning a total of 148.83 Mb. Most SVs were small, with 91.52% of insertions and 92.96% of deletions shorter than 1 kb, whereas only approximately 1% exceeded 10 kb in length (**Supplementary Tables 3 and 4**). A substantial proportion of SVs (36.33%) were associated with transposable elements (TEs), with DNA transposons containing terminal inverted repeats (DNA-TIRs) representing the most abundant class. Notably, 6.23% of SVs overlapped coding sequence (CDS) regions, and 24.33% of genes had CDSs affected by at least one SV (**Supplementary Table 5**).

To explore SV diversity at the population level, we performed deep genome resequencing of a core collection^20^ of 206 *C. pepo* accessions at an average depth of 53.4× (**Supplementary Table 6**). Misidentification and taxonomic inconsistency are well-recognized challenges in plant germplasm collections and are particularly pronounced in *C. pepo* due to its bewildering phenotypic variation. Among the 206 accessions, one was initially annotated as *C. pepo* subsp. *fraterna*, seven as *C. pepo* subsp. *ovifera* var. *ozarkana*, six as *C. pepo* subsp. *ovifera* var. *texana*, and two as *C. pepo* subsp. *pepo*, whereas the remaining 190 were broadly classified as *C. pepo* without further taxonomic resolution (**Supplementary Table 6**). To refine these assignments, we integrated phylogenetic relationships inferred from genome-wide SNPs (**Supplementary Fig. 3**) with manual inspection of fruit morphology. This framework enabled the confident classification of all 190 previously unresolved accessions into specific subspecies and/or cultivar groups. In addition, two accessions were identified as misclassified and reassigned to their correct taxonomic groups (**Supplementary Table 6**). After taxonomic correction, this global collection comprised 12 wild accessions representing three wild subspecies (1 *C. pepo* subsp. *fraterna*, 5 *C. pepo* subsp. *ovifera* var. *texana*, and 6 *C. pepo* subsp. *ovifera* var. *ozarkana*) and 194 cultivated accessions (33 *C. pepo* subsp. *ovifera* var. *ovifera* and 161 *C. pepo* subsp. *pepo*), including both landraces and modern cultivars. These accessions spanned five continents and a broad geographic range, from ∼37°S to 60°N latitude and from ∼120°W to 142°E longitude, encompassing tropical, subtropical, temperate, and continental climatic regimes (**Fig. 1c and Supplementary Table 6**).

Using the high-depth resequencing data, we genotyped SVs in the graph-based pangenome across all 206 accessions with PanGenie^21^. Comparison with assembly-derived SVs, as well as leave-one-out validation, supported the accuracy of SV genotyping (**Supplementary Fig. 1**). We observed marked variation in both the number and length of SVs among populations and individual accessions (**Fig. 1d, Supplementary Fig. 2 and Supplementary Table 7**). In addition, we identified 5,040,318 high-quality SNPs from the same resequencing dataset (**Supplementary Table 8**). Together, these SNPs and SVs constitute an integrated population-scale genomic resource for investigating the evolutionary history and population differentiation in *C. pepo*.

### Parallel domestication of two subspecies of *C. pepo*

The domestication of *C. pepo* is among the earliest known plant domestication events, with archaeological evidence indicating human use of *C. pepo* as food as early as 10,000 years before present^12^. Over millennia of cultivation and selection, *C. pepo* has developed remarkable morphological diversity and broad ecological adaptability^15^. Archaeological and molecular evidence indicates that, prehistorically, *C. pepo* was widely distributed in North America, from southern Mexico and perhaps Guatemala to northwards and eastwards through the eastern U.S.A. perhaps as far as the eastern states bordering Canada. The population of Southern North America, subsp. *pepo*, and the population of Northern North America, subsp. *ovifera*, underwent at least two independent domestication events, the former giving rise to pumpkins and the latter to squash^10,13,14^. However, the evolutionary relationships among wild relatives and cultivated lineages, as well as the extent to which these domestication events were truly independent, have remained unclear.

Phylogenetic analyses based on genome-wide variants have placed a third, undomesticated taxon, subsp. *fraterna*, as allied with subsp. *ovifera* (including var. *texana*, var. *ozarkana*, and var. *ovifera*) or sister to both of the other subspecies, with subsp. *ovifera* and subsp. *pepo* forming two major derived clades (**Supplementary Figs. 3** and **4**). Within the *ovifera* lineage, the cultivated form (*C. pepo* subsp. *ovifera* var. *ovifera*) clustered most closely with the wild taxon var. *texana*, supporting a potential direct domestication origin from a *texana*-like ancestor. In contrast, subsp. *pepo*, the globally distributed and economically dominant cultivated lineage, lacks an extant wild progenitor, obscuring its origin in simple phylogenetic reconstructions. Population structure analyses further supported a clear separation between the two major subspecies (**Supplementary Fig. 5**). At *K* = 4, subsp. *ovifera* and subsp. *pepo* formed distinct genetic clusters, while the wild relatives within the *ovifera* lineage (var. *texana* and var. *ozarkana*) showed finer-scale differentiation from the cultivated *ovifera* group. Notably, Southern North American accessions of subsp. *pepo* formed a partially distinct cluster (**Fig. 2a** and **Supplementary Fig. 5**). Similar patterns were observed in principal component analysis (PCA) and genome-wide genetic differentiation (*F*_ST_). PC1 (38.35%) clearly separated the two subspecies, whereas PC2 (3.03%) captured within-subspecies variation, including differentiation among varieties and geographic groups (**Fig. 2b**). Consistently, *F*_ST_ values were low within subspecies but markedly elevated between them (**Fig. 2c**), indicating strong divergence between the two domesticated gene pools.

**Figure 2.**
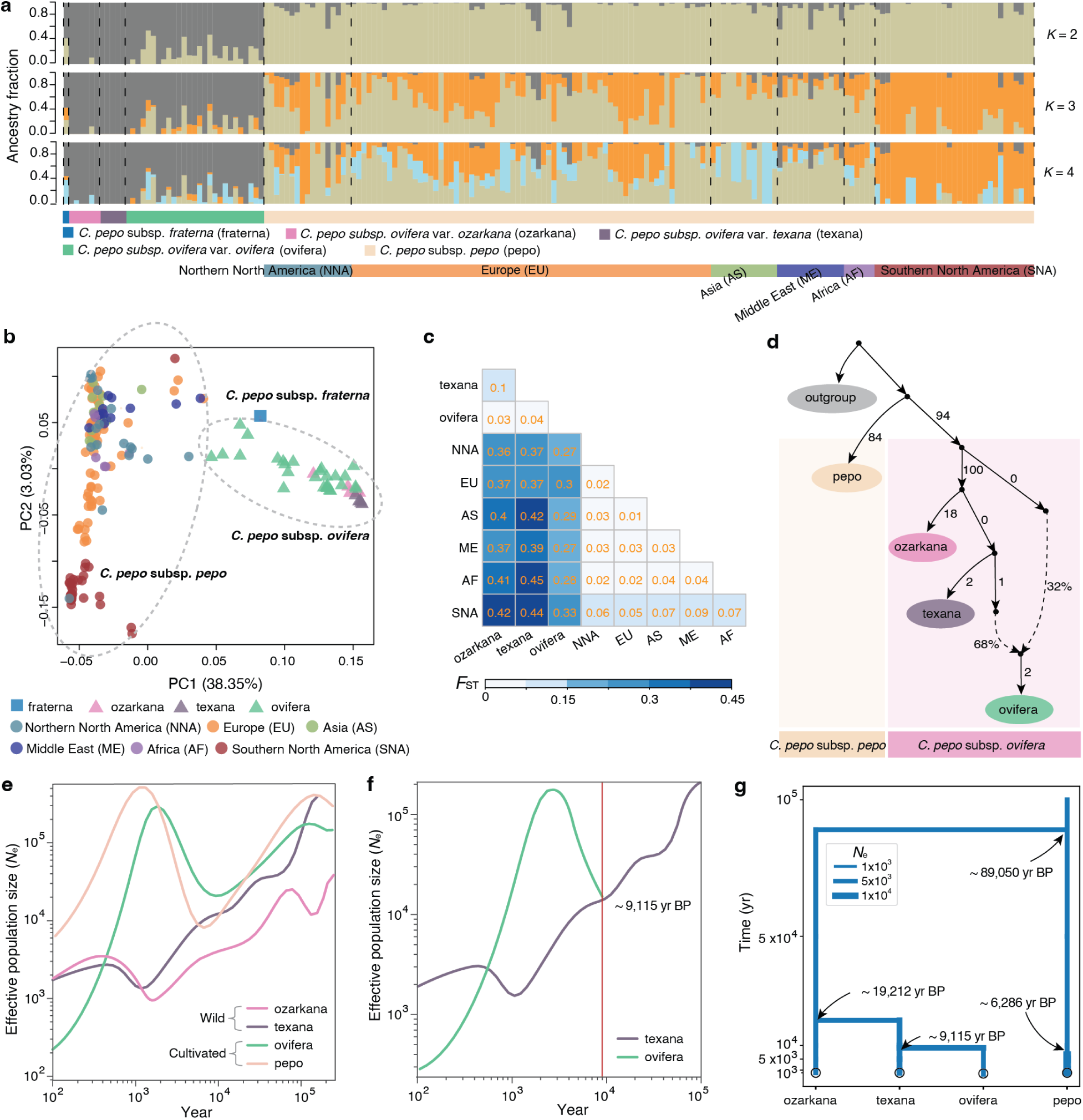
Genetic structure, divergence, and demographic history in *C. pepo*. **a,** Population structure inferred at *K* = 2–4. Each bar represents an accession and is colored according to ancestry proportion. **b,** Principal component analysis (PCA) of *C. pepo* accessions. **c,** Pairwise fixation index (*F*_ST_) among populations. **d**, Inferred admixture graph showing relationships among *C. pepo* groups. Numbers on branches indicate drift parameters, and dashed arrows indicate admixture events. **e**, Changes in effective population size (*N_e_*) over time in *C. pepo* groups, inferred using SMC++. **f,** Divergence time between cultivated *C. pepo* subsp. *ovifera* var. *ovifera* and its closest wild relative *C. pepo* subsp. *ovifera* var. *texana*, estimated using SMC++. **g**, Schematic representation of divergence times among major *C. pepo* groups. Branch widths represent branch-specific effective population sizes (*N_e_*) estimated using momi2.

Although these patterns revealed substantial divergence between subsp. *pepo* and subsp. *ovifera*, they did not by themselves establish whether the two cultivated lineages originated independently. To address this question, we evaluated alternative evolutionary scenarios using admixture graph models implemented in qpGraph^22^. Models assuming a single domestication origin–either deriving both cultivated lineages from a *texana*-like ancestor or treating subsp. *pepo* as derived from cultivated *ovifera*–were strongly rejected based on large absolute Z-scores (**Supplementary Figs. 6a,b**). In contrast, models incorporating an ancestral lineage not represented by the extant wild populations sampled in this study showed a substantially improved fit, supporting an independent origin of the subsp. *pepo* lineage (**Supplementary Fig. 6c**). We subsequently performed an unconstrained heuristic search of alternative graph topologies. Among the 92 candidate models evaluated, three showed acceptable fits (|Z| < 3; **Supplementary Fig. 6d– f**), all indicating an early separation of the lineage leading to *subsp. pepo* from that containing *texana*, *ozarkana*, and cultivated *ovifera*. The best-fitting model (|Z| = 2.057; **Fig. 2d and Supplementary Fig. 6d**) placed subsp. *pepo* as an early-diverging lineage relative to the *ovifera* clade, whereas cultivated *ovifera* was derived primarily from a *texana*-like ancestor with a substantial contribution from an additional ancestral lineage. Together, these results reject a single-origin model and indicate that the two cultivated subspecies arose independently from distinct ancestral gene pools.

Demographic inference further supported this scenario. Both cultivated lineages (*pepo* and *ovifera*) underwent pronounced population bottlenecks between ∼5,000 and 10,000 years ago, followed by population expansion associated with human cultivation and dispersal (**Fig. 2e**). In contrast, wild populations (*ozarkana* and *texana*) showed a gradual decline in effective population size over time (**Fig. 2e**). Integrating SMC++ ^23^ and momi2 ^24^ analyses, we reconstructed a temporal framework for the diversification of *C. pepo* (**Fig. 2f,g**). The two major subspecies were estimated to have diverged 89,050 years before present, followed by the split between var. *texana* and var. *ozarkana* 19,212 years before present. The *ovifera* lineage was inferred to have been domesticated 9,115 years ago from a *texana*-like ancestor (**Fig. 2f**), coinciding with the onset of agriculture in North America^25^. For subsp. *pepo*, we inferred a pronounced demographic bottleneck approximately 6,286 years ago. This bottleneck is more recent than archaeological evidence for domestication^10^ (∼8,000–10,000 years ago), suggesting that the inferred bottleneck may reflect a later population contraction associated with domestication rather than the initial onset of domestication. Historical gene flow involving subsp. *pepo* and unsampled ancestral or related lineages, together with incomplete sampling of ancestral diversity, may also have influenced the inferred timing of this event. Pairwise two-dimensional site-frequency spectra (2D SFS) generated by msprime^26^ simulations under the momi2 ^24^ demographic model broadly recapitulated the observed spectra, supporting the overall adequacy of the inferred demographic model (**Supplementary Fig. 7**). Together, these results support a model of parallel domestication in *C. pepo*, in which two deeply diverged ancestral gene pools independently gave rise to distinct cultivated lineages.

### Differential selective constraints on SNPs and SVs in *C. pepo*

To characterize the evolutionary forces shaping genomic variation in *C. pepo*, we investigated the contributions of both SNPs and SVs to adaptive variation across the wild populations and across each of the two cultivated subspecies, *ovifera* and *pepo*. We estimated the distribution of fitness effects (DFE) from population allele frequency spectra using polyDFE^27^, with synonymous SNPs as the neutral reference. We observed that in all three populations, SVs experienced stronger purifying selection than SNPs, with a larger proportion of SVs showing strongly deleterious effects (*N*_e_*S* < -100) (**Supplementary Fig. 8a-c**). This pattern is consistent with the larger functional impacts of SVs, which frequently disrupt coding or regulatory regions and are therefore rapidly purged by selection, as also reported in cucumber^28^.

Despite pervasive purifying selection, a small subset of SVs persisted at moderate to high frequencies, suggesting that these variants are not subject to strong purifying selection and may include neutral, nearly neutral, or potentially advantageous variants. To further quantify the adaptive variation, we estimated the proportion of fixed adaptive mutations (α) and found higher values for SNPs (∼0.40–0.45) than for SVs (∼0.19–0.21), consistent with the stronger purifying selection acting on SVs (**Supplementary Fig. 8d**). Therefore, incorporating SVs into population genomic frameworks provides a more comprehensive understanding of the evolutionary forces that have shaped the genomic architecture and adaptive potential of *C. pepo*.

### Climate-driven genomic differentiation in *C. pepo*

Accessions in the *C. pepo* core collection span a broad geographic range and encompass diverse climatic backgrounds. Moreover, most cultivated accessions represent landraces and historical or older varieties, with many collected before the mid-1980s (**Supplementary Table 6**). As a result, these accessions have likely been maintained locally for extended periods, enabling them to retain genetic signatures associated with local environmental conditions. Together, this core collection provides a valuable framework for investigating the role of climate in shaping genomic variation across *C. pepo* populations. Accordingly, as in previous studies^29–33^, we integrated both wild and cultivated accessions to identify genomic variants associated with environmental gradients across the species’ range.

We performed redundancy analysis (RDA)^34^ using 19 bioclimatic variables, including 11 temperature-related (BIO1–BIO11) and 8 precipitation-related variables (BIO12–BIO19) (**Supplementary Table 9**). Both SNP- and SV-based RDAs revealed strong associations between genomic variation and multiple climatic predictors, with the first two RDA axes explaining substantial proportions of total genomic variance (**Fig. 3a,b**). The full RDA models were highly significant for both SNP and SV datasets (*P* < 0.001). Because environmental variables are often spatially structured, we further assessed the robustness of these climate–genome associations by accounting for spatial autocorrelation using distance-based Moran’s eigenvector maps (PCNM)^35^. After incorporating spatial structure, climatic predictors remained highly significant (*P* < 0.001), demonstrating that climate significantly contributes to genomic differentiation in *C. pepo*.

**Figure 3.**
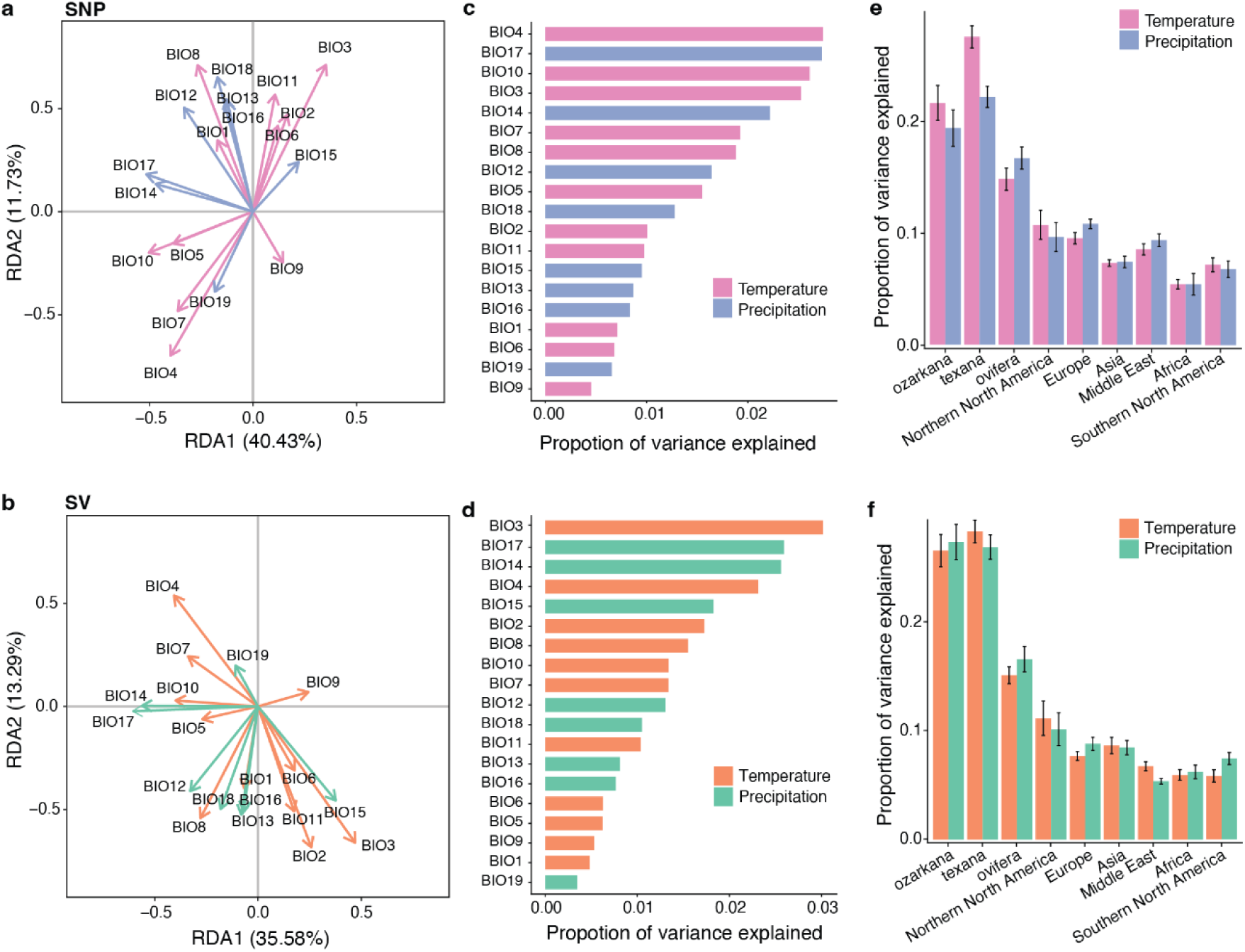
Climate-driven genomic differentiation in *C. pepo*. **a-b**, Contribution of bioclimatic variables (BIO1–BIO19) to the first two principal components derived from redundancy analysis (RDA) based on SNPs (**a**) and SVs (**b**). **c-d,** Proportion of genomic variance explained by individual bioclimatic variables for SNPs **(c)** and SVs **(d)**. **e-f,** Climate-associated genomic variance partitioned across subspecies and geographic regions for SNPs (**e**) and SVs (**f**).

Among climatic predictors, temperature-related variables explained more genomic variation than precipitation-related variables. In particular, BIO3 (isothermality) and BIO4 (temperature seasonality) showed the greatest explanatory power, highlighting that temperature gradients contribute more strongly to genomic differentiation in *C. pepo* (**Fig. 3c,d**). Notably, climatic factors explained a larger proportion of genetic variation in wild populations than in cultivated populations (**Fig. 3e,f**). This pattern likely reflects the longer evolutionary exposure of wild relatives to diverse environmental conditions, as well as relaxed environmental selection under cultivated conditions.

### Genomic loci associated with climatic gradients

Given the strong contribution of climatic gradients to genome-wide differentiation, we next sought to identify specific genomic loci associated with climatic variation using genome–environment association (GEA) analyses. Because the 19 bioclimatic variables related to temperature and precipitation are highly correlated (**Supplementary Fig. 9**), we reduced dimensionality by performing separate principal component analyses (PCAs) for temperature-related and precipitation-related variables. The first two principal components explained 76.28% and 83.32% of the total variance for temperature and precipitation, respectively, and were retained as composite climatic predictors for downstream analyses (**Supplementary Fig. 10**).

Using these major climatic principal components, we identified climate-associated loci with Bayenv2 ^36^. Both SNPs and SVs exhibited abundant associations with environmental variables **(Fig. 4a,b** and **Supplementary Fig. 11a,b**), with effect sizes exceeding neutral expectations based on permutation analyses, indicating robust environmental signals beyond neutral demographic structure. We identified 13,782 climate-associated SNPs and 7,609 SVs, which were broadly distributed across the genome rather than clustered in a few genomic regions, suggesting widespread environmentally driven differentiation. This dispersed pattern supports a largely polygenic architecture of climatic adaptation in *C. pepo*, likely involving many loci of small to intermediate effect rather than a limited set of major-effect variants.

**Figure 4.**
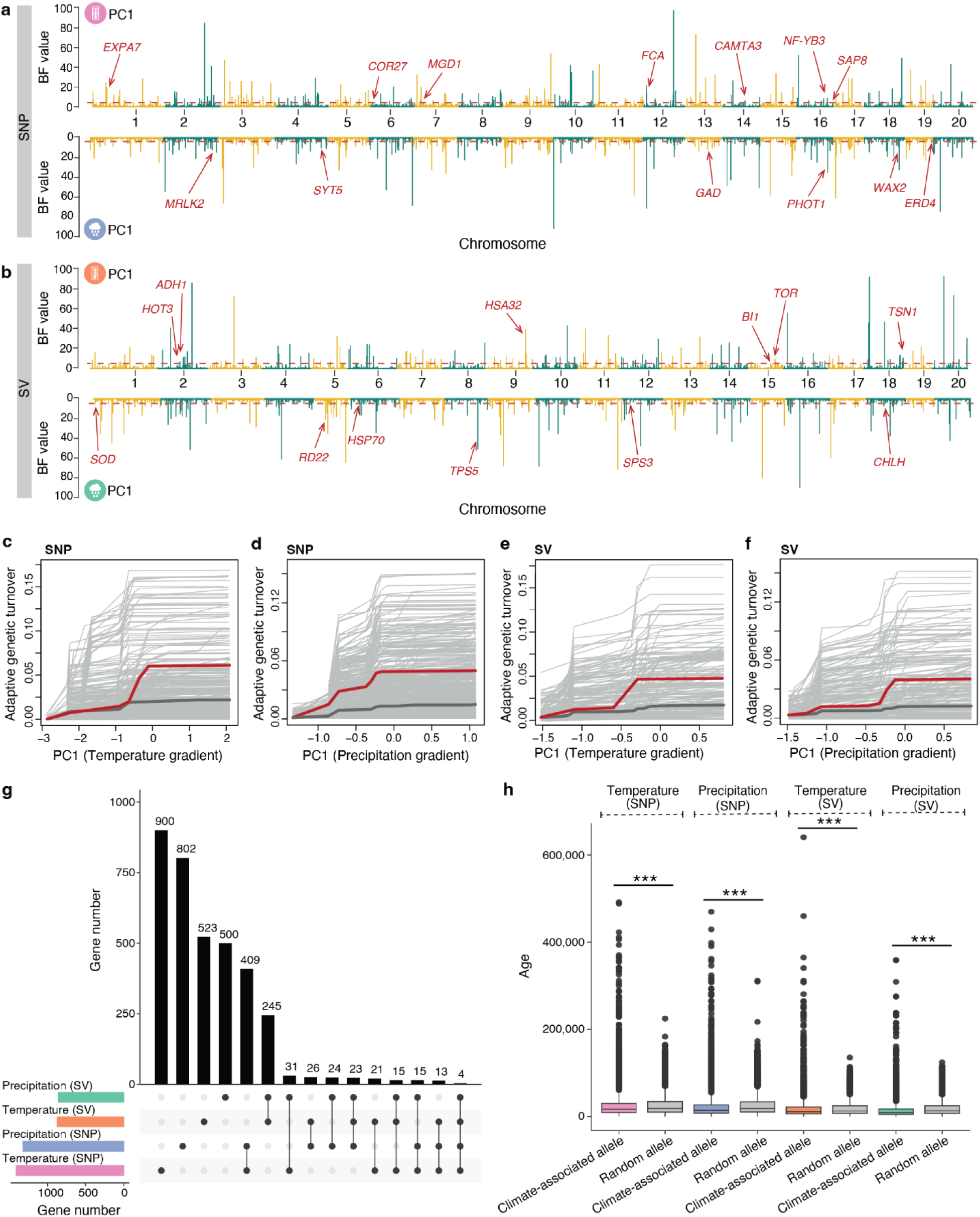
Climate-associated adaptive genetic variation inferred from SNPs and SVs. **a,b**, Genome-wide distribution of climate-associated SNPs (**a**) and SVs (**b**) along PC1 of temperature (top panels) and precipitation (bottom panels) gradients. The dashed red line indicates the significance threshold, and representative candidate genes near the top signals are shown. Details of the representative candidate genes are provided in **Supplementary Table 12**. **c–f,** Turnover of adaptive SNPs (**c**,**d**) and SVs (**e**,**f**) along PC1 of temperature and precipitation gradients. Thin light-grey lines represent turnover trajectories of individual loci, thick dark-grey lines indicate the mean cumulative turnover of random loci, and colored thick lines denote the mean turnover of climate-associated loci. **g,** UpSet plot showing overlap among candidate genes associated with climate adaptation. **h,** Estimated variant ages of temperature- and precipitation-associated alleles and random alleles. Asterisks indicate significant differences between climate-associated and random variants (*p* < 0.001, Wilcoxon rank-sum test). For each boxplot, the lower and upper bounds indicate the first and third quartiles, respectively, the center line indicates the median, and the whiskers extend to 1.5× the interquartile range.

To characterize how the climate-associated loci identified by Bayenv2 vary along environmental gradients, we applied Gradient Forest models^37^ to quantify nonlinear allele-frequency turnover. Both adaptive SNPs and SVs exhibited pronounced nonlinear allele turnover across climatic gradients, with significantly higher turnover rates than randomly sampled loci (**Fig. 4c–f** and **Supplementary Fig. 11c–f**). Temperature-related variables produced stronger genetic turnover than precipitation-related variables, consistent with temperature explaining a larger fraction of genomic variation (**Fig. 3c,d**). The consistent turnover patterns observed for SNPs and SVs further support their associations with climatic variation and highlight distinct genomic responses along environmental gradients in *C. pepo*.

In total, we identified 3,551 genes associated with climatic gradients, spanning functional categories related to genome maintenance, transcriptional regulation, cell wall biogenesis, and metabolic and transport processes (**Supplementary Tables 10-11** and **Supplementary Fig. 12**). Among these, a subset had orthologs previously implicated in abiotic-stress responses or related physiological processes in *C. pepo* or other plant species (**Supplementary Table 12**). For example, the candidate gene *RD22* expression was upregulated in melon leaves after 7–21 days of water deficit^38^. The *EXPA7* candidate belongs to the expansin family, several members of which show abiotic-stress-responsive expression in *C. pepo*^39^ (**Fig. 4a,b**). These observations provide supporting evidence for the potential involvement of these gene families in environmental responses, although the specific loci identified here remain to be functionally validated. Notably, most climate-associated genes were specific to particular variant-environment combinations, whereas only a small subset was shared between SNP- and SV-based signals or between temperature- and precipitation-associated loci (**Fig. 4g**). SNPs and SVs therefore capture largely complementary sets of climate-responsive genes, highlighting their distinct contributions to climate adaptation.

We then analyzed the ages of climate-associated variants and found that these variants were generally younger than genome-wide background variants for both SNPs and SVs (**Fig. 4h**). Although absolute allele-age estimates can be influenced by demographic and genomic factors, including effective population size, mutation and recombination rates, and allele frequency^40,41^, additional sensitivity analyses showed that this younger age pattern persisted after accounting for genomic location and allele-frequency differences, and that more widely shared climate-associated alleles tended to be older (**Supplementary Figs. 13–16**). Simulation-based validation showed a strong correspondence (*r* = 0.868) between estimated and true allele ages (**Supplementary Fig. 17**), supporting the reliability of the age estimates. Together, these patterns suggest that recent environmental pressures have strongly shaped the adaptive landscape of *C. pepo*, particularly through the retention of recently arisen, functionally impactful variants.

### Predicting climate vulnerability in *C. pepo*

To forecast how *C. pepo* populations may respond to future climates, we integrated ecological niche modeling with genetic offset analyses informed by climate-associated SNPs and SVs. Species distribution models (SDMs)^42^ projected substantial shifts in suitable habitat under both mid-century (2040–2060) and late-century (2080–2100) climate scenarios (**Fig. 5a,b**). The wild relatives *C. pepo* subsp. *ovifera* var. *ozarkana* and *var. texana* were projected to experience pronounced range contractions accompanied by northward shifts, particularly across the southern United States, with range reductions of 65.46% and 89.35% by mid-century and 92.34% and 94.26% by late-century, respectively (**Fig. 5a**, **Supplementary Fig. 18** and **Supplementary Table 13**). In contrast, the two cultivated taxa (var. *ovifera* and subsp. *pepo*) displayed more moderate overall contractions, together with projected expansions into higher latitudes in both hemispheres and contractions in warmer regions such as Central America and northern Africa (**Fig. 5a,b, Supplementary Figs. 18 and 19** and **Supplementary Table 13**). Together, these projections indicate that future climatic regimes may substantially reshape the geographical distribution of suitable habitat for *C. pepo*, underscoring the need for conservation strategies that protect its full diversity, particularly wild relatives that harbor unique reservoirs of adaptive genetic variation.

**Figure 5.**
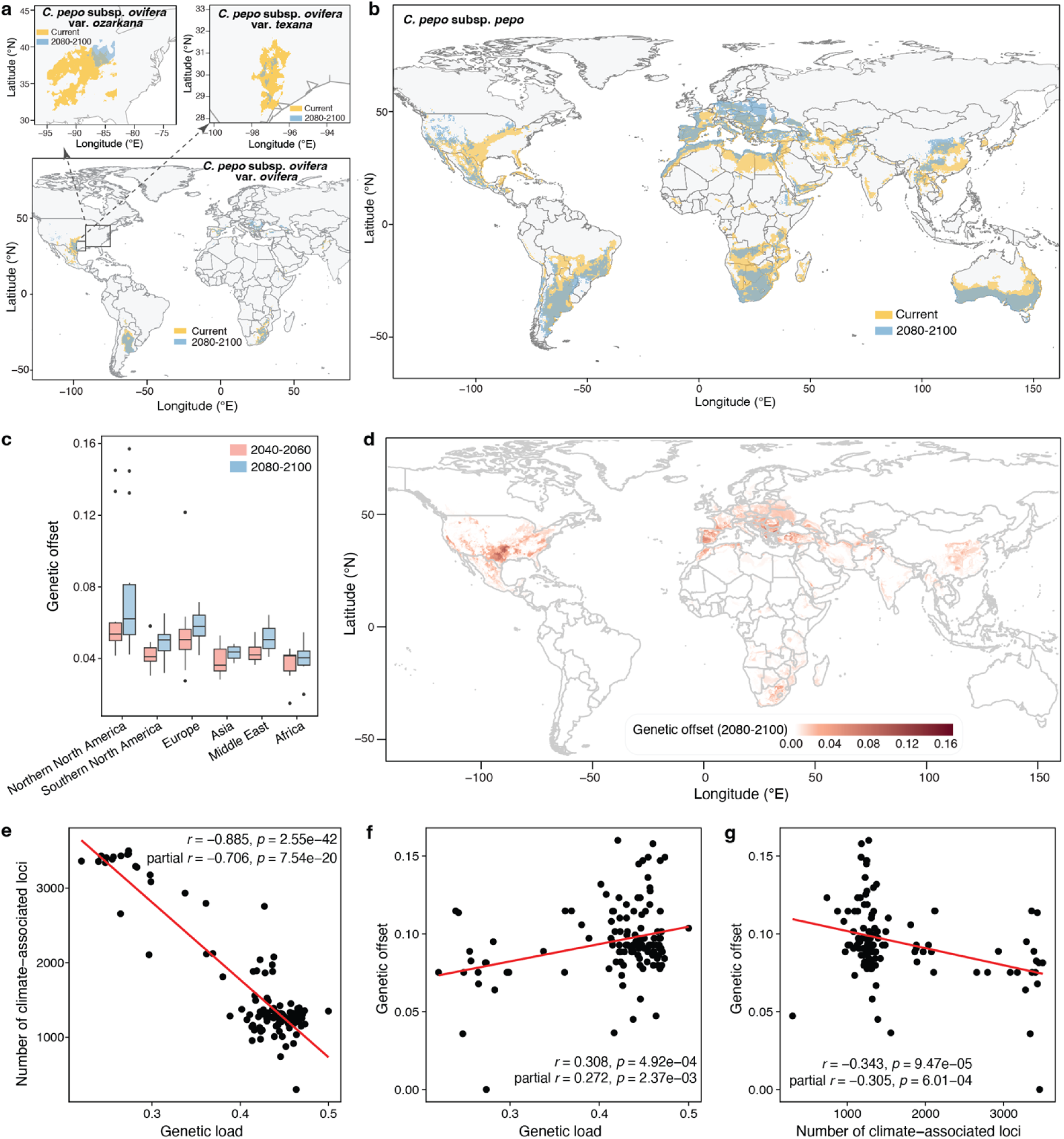
Projected range shifts and genomic vulnerability for *C. pepo* under future climate scenarios. **a**, Predicted current and late-century (2080–2100) climatically suitable areas for wild *C. pepo* subsp. *ovifera* var. *ozarkana* and var. *texana* (top panels) and cultivated *C. pepo* subsp. *ovifera* var. *ovifera* at the global scale (bottom panel), inferred from species distribution models. Boxes in the bottom panel indicate Northern North America regions shown in the top panels. **b**, Predicted current and late-century (2080–2100) climatically suitable areas for *C. pepo* subsp. *pepo* at the global scale. **c**, Regional comparison of genetic offset across continents under mid-century (2040–2060) and late-century climate scenarios. For each boxplot, the lower and upper bounds indicate the first and third quartiles, respectively, the center line indicates the median, and the whiskers extend to 1.5× the interquartile range. **d**, Spatial distribution of genetic offset projected for late century for the entire *C. pepo* population. **e–g**, Pearson correlations between genetic load, number of climate-associated loci, and genetic offset. Each point represents an individual accession. Maps in **a, b** and **d** were created using the R package rworldmap (https://cran.r-project.org/web/packages/rworldmap/index.html).

To incorporate evolutionary potential into assessments of future maladaptation risk, we estimated genetic offset, which quantifies the magnitude of allele-frequency shifts required for populations to maintain their current genotype–environment relationships under projected future climates. Because SNPs and SVs capture complementary components of climate-associated genomic variation (**Fig. 4g**), both variant classes were jointly integrated into the genomic prediction framework. In the combined SNP and SV analysis, genetic offset increased from mid-century to late-century climate scenarios, indicating a progressively rising risk of maladaptation over time (**Fig. 5c**). Separate SNP- and SV-based analyses revealed broadly consistent regional patterns, although the magnitude of genetic offset differed between the two variant classes (**Supplementary Fig. 20**). Notably, populations from Northern North America exhibited the highest genetic offset values, suggesting that they would require substantial allele-frequency changes to maintain local adaptation under future climatic conditions (**Fig. 5d** and **Supplementary Fig. 21**).

To investigate how intrinsic genomic constraints may shape vulnerability to future climates, we examined relationships among genetic load, the number of climate-associated loci, and genetic offset (**Fig. 5e–g**). Genetic load, estimated as the relative burden of derived nonsynonymous SNPs and SVs, was negatively correlated with the number of climate-associated loci (*r* = -0.885, *P* = 2.55 × 10^-42^; **Fig. 5e**). This association was attenuated but remained significant after controlling for effective population size (*N*_e_) (partial *r* = -0.706, *P* = 7.54 × 10^-20^) (**Supplementary Fig. 22**). Genetic load was also positively associated with genetic offset (*r* = 0.308, *P* = 4.92 × 10^-4^; partial *r* = 0.272, *P* = 2.37 × 10^-3^; **Fig. 5f**), suggesting that populations with elevated genetic load tended to show greater projected genomic changes under future climates. Conversely, the number of climate-associated loci was negatively correlated with genetic offset (*r* = -0.343, *P* = 9.47 × 10^-5^; partial *r* = -0.305, *P* = 6.01 × 10^-4^; **Fig. 5g**). We next compared genetic load and genetic offset with projected range change across taxonomic and geographic groups (**Supplementary Table 14**). The wild relatives *C. pepo* subsp. *ovifera* var. *ozarkana* and var. *texana* showed the greatest projected range contractions but relatively low genetic load and moderate genetic offset. In contrast, cultivated *C. pepo* groups generally showed less severe range contractions but higher genetic load, with some groups also exhibiting relatively high genetic offset. These contrasting patterns suggest distinct dimensions of climate vulnerability: wild populations may be more strongly affected by projected range contraction, whereas cultivated groups may face greater genomic constraints on adaptation. Together, these results highlight the interplay between ecological exposure and genomic adaptive capacity in shaping future climate vulnerability in *C. pepo* and underscore the need for conservation strategies tailored to the distinct vulnerabilities of individual populations.

## Discussion

Environmental variation poses a major challenge to both biodiversity conservation and agricultural sustainability^1^. Understanding how genetic variation underlies responses of species to environmental heterogeneity is therefore key to predicting evolutionary persistence and informing breeding strategies. By integrating genome-wide variation with climate-associated differentiation and predictive modeling, our study provides a comprehensive understanding of how *C. pepo* has evolved, adapted, and may respond to changing environmental conditions.

Our integrative analyses provide strong evidence for parallel domestication in *C. pepo*, in which two deeply diverged gene pools independently gave rise to the cultivated *pepo* and *ovifera* lineages. The two subspecies diverged approximately 89,000 years ago, followed by the split of *texana* and *ozarkana* around 19,000 years ago. Domestication of *ovifera* occurred ∼9,100 years ago from a texana-like ancestor, whereas *pepo* was domesticated likely from an unsampled or extinct progenitor. Despite their independent origins, both lineages experienced similar demographic trajectories, including domestication bottlenecks followed by expansion associated with human cultivation and dispersal. As these lineages expanded across diverse environments, climatic gradients emerge as an important force shaping genomic differentiation. Genome– environment association analyses identified numerous loci correlated with climatic principal components, broadly distributed across the genome and indicative of highly polygenic climatic adaptation. Gradient Forest modeling further reveals nonlinear allele turnover along temperature and precipitation gradients, demonstrating that climate-associated loci respond more strongly to changing environmental conditions than randomized variant sets. Importantly, SNPs and SVs capture complementary dimensions of this environmental responsiveness. Temporal analyses provided additional insight into the dynamics of adaptation: climate-associated variants are consistently younger than genome-wide background variants, suggesting an important contribution of relatively recent genetic variation to climate adaptation in *C. pepo*. Such variation may have arisen or increased in frequency during post-domestication expansion or in response to Holocene environmental variability. Although our GEA analyses incorporated both cultivated and wild accessions, the limited number of wild accessions in the current dataset precluded a robust wild-only analysis. Moreover, the historical movement of some accessions cannot be reconstructed from the available data, and human-mediated translocation may have contributed to some genome– environment associations. Expanded sampling of wild populations will therefore be important for independently validating climate-associated loci and further resolving patterns of local adaptation in *C. pepo*.

Projections under future climate scenarios indicate increasing genetic offset, particularly in Northern North American populations, along with continued range contraction of wild relatives, highlighting potential challenges for maintaining local adaptation under rapid environmental change. In addition to environmental drivers, intrinsic genomic properties may also influence adaptive potential. Our results suggest an association between genetic load and adaptative capacity, with populations harboring higher mutational burdens exhibiting fewer climate-associated loci and greater predicted genetic offset. This pattern emphasizes the buffering role of standing adaptive variation and reveals an evolutionary link among genetic diversity, mutational load, and adaptive potential. Such constraints are especially relevant for understanding why some populations are disproportionately vulnerable to future climates, even under similar environmental pressures.

Beyond providing insights into the evolutionary history and climatic adaptation of *C. pepo*, our study highlights the value of integrating multiple layers of genomic variation into predictive frameworks of environmental response. Most population genomic studies of crop adaptation have focused primarily on SNPs^31,43^, whereas our results demonstrate that SVs capture complementary patterns of climate-associated genomic variation. By jointly analyzing SNPs and SVs within a graph-based pangenome framework, we provide a more comprehensive view of the genomic architecture associated with environmental adaptation. Nonetheless, the limited number of genome assemblies currently available may not capture the full spectrum of SV diversity in *C. pepo*. In particular, the absence of wild assemblies from the graph pangenome may have led to underrepresentation of wild-specific SVs, potentially contributing to the observed differences in the inferred DFE between wild and cultivated populations. Incorporating additional high-quality genome assemblies, particularly from wild populations, will provide a more comprehensive representation of SV diversity and enable more robust comparisons across populations. Such integrative approaches will be increasingly important for understanding how complex genomes respond to environmental changes and for identifying genetic variants that may contribute to adaptation under future environmental conditions. More broadly, our findings emphasize that adaptive potential emerges from the interplay between beneficial and deleterious variation. While standing adaptive variation provides the raw material for evolutionary responses to changing environments, the accumulation of deleterious mutations can simultaneously constrain this process. The strong associations observed among genetic load, adaptive variation, and predicted genetic offset highlight the importance of considering both evolutionary opportunities and genomic constraints when evaluating adaptive potential in crop populations.

Overall, our findings reveal how the interplay among selection, genomic variation, and genetic constraints shapes the adaptive trajectory of *C. pepo* across temporal and environmental scales. Conserving genetically diverse and locally adapted populations will be critical for sustaining its evolutionary potential. This integrative genomic framework also offers a broadly applicable approach for assessing adaptive capacity and guiding conservation and breeding strategies in other crops.

## Methods

### Genome sequencing, assembly and annotation

High-molecular-weight genomic DNA was extracted from young fresh leaves using the cetyltrimethylammonium bromide (CTAB) method^44^. SMRTbell libraries were constructed following the standard protocol provided by PacBio and sequenced on a PacBio Sequel II platform using the circular consensus sequencing mode to generate HiFi reads. For all accessions, HiFi reads were processed using HiFiAdapterFilt^45^ (v2.0.1) to remove adapter sequences. The cleaned HiFi reads were assembled into contigs using hifiasm^46^ (v0.14.2-r315). Shorter contigs that were represented by a longer contig with at least 99% identity and coverage were identified and removed as haplotypic duplications according to BLAST+ ^47^ (v2.12.0+) alignments. Potential contaminant sequences from microorganisms and organelle genomes were also identified and removed by comparing contig sequences against the NCBI nt database using BLAST+ ^47^ (v2.12.0+). For the accession C39, cleaned contigs were anchored to chromosomes by integrating three published genetic maps^48–50^, a Hi-C dataset from DNA Zoo using the 3D-DNA pipeline^51^ (v180922), as well as the genome assembly of *C. pepo* accession MU-CU-16 ^52^ for reference-guided anchoring using RagTag^53^ (v2.0.1). For the remaining eight accessions (C31, C38, JOL, OVF, SPR, TRF, 114a, and VSP), chromosome-level assemblies were generated using RagTag^53^ (v2.0.1) with the C39 genome as reference.

Repeat sequences were first identified and classified in each of the nine *C. pepo* genome assemblies using EDTA^54^ (v2.1.3) to generate genome-specific de novo TE libraries. The resulting libraries were then combined and filtered to remove redundant TE sequences, yielding a non-redundant pangenome TE library. This library was used as a custom repeat library in RepeatMasker^55^ (v4.1.1) to identify and annotate repetitive sequences across all nine *C. pepo* genome assemblies. Protein-coding genes were initially predicted from the repeat-masked genomes using the MAKER pipeline^56^ (v3.01.03), which integrates *ab initio* predictions, transcript evidence, and homologous protein evidence. Specifically, AUGUSTUS^57^ (v3.4.0) and SNAP^58^ (v2006-07-28) were used for *ab initio* gene prediction, and RNA-seq data from various tissues downloaded from NCBI (accessions PRJNA239659, PRJNA752874, PRJNA339848, PRJNA386743, PRJNA437911, and PRJNA439198) were used for transcript evidence. For protein homology prediction, protein sequences from Arabidopsis (TAIR10), melon (*Cucumis melo*)^59^, cucumber (*Cucumis sativus*)^60^, watermelon (*Citrullus lanatus*)^61^, wax gourd (*Benincasa hispida*)^62^, snake gourd (*Trichosanthes anguina*)^63^, and chayote (*Sechium edule*)^64^, *Cucurbita pepo*^52^, *Cucurbita maxima*^65^, *Cucurbita moschata*^65^, and the UniProt (Swiss-Prot plant division) database were aligned to the genomes using Spaln^66^ (v3.0.1). Predicted gene models were subsequently transferred across assemblies using Liftoff^67^ (v1.5.1). Furthermore, Helixer (v0.3.2), a gene prediction tool for eukaryotes that combines deep learning and a hidden Markov model^68^, was also applied to all nine assemblies. Finally, results from all approaches were integrated using EVidenceModeler^69^ (v1.1.1) to generate the final set of predicted protein-coding genes for each genome assembly.

### Gene-based and graph-based pangenome construction

Protein sequences from the nine *C. pepo* genomes were clustered into gene families (pan-gene clusters) using OrthoFinder^70^ (v2.5.5).

The graph-based pangenome was constructed using the Minigraph-Cactus package^71^ with the nine reference-level genome assemblies. Pangenome construction began with the C39 genome as the initial graph, to which the remaining eight assemblies were sequentially mapped. The resulting genome graph was indexed and exported to VCF format using the vg toolkit^72^ and filtered using vcfbub (https://github.com/pangenome/vcfbub) with parameters ‘-l 0 -r 10000’.

### Short-read sequencing, SNP calling, and SV genotyping and validation

Genomic DNA was isolated from young fresh leaf tissues using the Omega Mag-Bind Plant DNA DS Kit (M1130, Omega Bio-Tek, Norcross, GA) following the manufacturer’s instructions. Shotgun libraries were constructed from the extracted DNA and sequenced on the BGISEQ-500 platform to generate 150-bp paired-end reads. Raw sequencing reads were processed to remove adaptors and low-quality sequences using Trimmomatic^73^ (v0.39). For SNP calling, cleaned reads were aligned to the C39 genome using BWA-MEM^74^ (v0.7.17-r1188) with default parameters, and duplicate read pairs were marked. SNPs were called using the Sentieon package (https://www.sentieon.com/) and filtered using GATK^75^ (v4.2.5.0) with parameters ‘QD < 2.0 || FS > 60.0 || MQ < 40.0 || MQRankSum < -12.5 || ReadPosRankSum < -8.0’. Furthermore, only biallelic SNPs with minor alleles present in at least two accessions were retained.

SVs in the graph-based pangenome were genotyped across the resequenced *C. pepo* accessions with cleaned resequencing reads using PanGenie^21^ (v3.0.2) and default parameters. An SV was considered TE-associated when at least 50% of its genomic interval overlapped an annotated TE. SNPs and SVs were functionally annotated using SnpEff^76^ (v5.1).

We evaluated PanGenie SV genotyping accuracy against assembly-derived SV calls using two different approaches. First, for accessions with genome assemblies, we compared PanGenie genotypes with the corresponding assembly-derived genotypes and calculated precision, recall, F1 score, and accuracy. We also assessed concordance between SV allele frequencies estimated by the two approaches using Pearson correlation. Second, we performed leave-one-out validation for the eight *C. pepo* accessions with both genome assemblies and short-read sequencing data. For each accession, we reconstructed the graph excluding its assembly, genotyped its short reads with PanGenie, and compared the resulting SV genotypes with those derived from its assembly.

### Population genomic analyses

Maximum-likelihood phylogenetic trees were constructed separately with the complete set of SVs and 109,659 transversion SNPs at fourfold degenerate sites (4DTv) using RAxML^77^ (v8.2.13) under the GTRGAMMA model with 100 bootstrap replicates. Three watermelon accessions^78^ (GenBank accession numbers: SRR8751845, SRR8751846, and SRR8751851) were used as the outgroup. Phylogenetic trees were visualized using iTOL^79^ (v7). Population structure was inferred using ADMIXTURE^80^ (v1.3.0) with SNPs having a minor allele frequency (MAF) ≥ 0.05 and linkage disequilibrium (LD) ≤ 0.2. Principal component analysis (PCA) was performed using PLINK^81^ (v1.9). Populations fixation index (*F*_ST_) was calculated separately with SNPs and SVs using VCFtools^82^ (v0.1.15) in 100-kb sliding windows with a 10-kb step size.

### Reconstruction of demographic history

Effective population sizes were inferred using SMC++ ^23^ (v1.15.4.dev18+gca077da). The genome of each accession was treated as a single haplotype, with occasional heterozygous sites resolved by randomly selecting one allele. For each population, haplotypes from different accessions were combined to create pseudo-diploid genotypes. A total of 15 pseudo-diploid genotypes were randomly selected for analysis. A mutation rate of 6.5 × 10⁻⁹ per site per generation^83^ and a generation time of 1 year were applied. The SMC++ analysis was repeated 20 times for each population, with pseudo-diploid genotypes resampled in each iteration.

Admixture graph analyses were performed using AdmixTools^22^. We first evaluated predefined graph topologies representing alternative domestication scenarios using qpGraph^22^ and then explored alternative graph topologies using the heuristic search implemented in qpBrute^84^. During the heuristic search, only the outgroup was specified; relationships among ingroup populations and the number and placement of admixture events were allowed to vary. A total of 92 candidate models were generated and evaluated, with models having |Z| < 3 considered acceptable. Among the retained models, the best-fitting model was identified using the admixturegraph package^85^ based on marginal likelihoods and Bayes factors. No additional SNP subsampling was performed for the admixture graph analyses, and an outgroup was included to root the admixture graphs. qpGraph was run with a block size of 0.05 Morgans for block-jackknife estimation and with the parameter ‘useallsnps: YES’.

Demographic parameters were further estimated using momi2 ^24^ (v2.1.19). Folded site-frequency spectra (SFS) were computed and divided into 100 blocks for jackknife and bootstrap resampling. The demographic topology was predefined based on the major population relationships supported by the phylogenetic and admixture graph analyses. The split between *texana* and cultivated *ovifera* was fixed at 9,115 years ago based on the independent SMC++ estimate, whereas the remaining demographic parameters were estimated by momi2 within this simplified branching framework. Because no extant wild population represents the direct progenitor of subsp. *pepo*, an equivalent wild–cultivated split time could not be estimated using SMC++. However, the SMC++ demographic trajectory for subsp. *pepo* showed a pronounced reduction in effective population size (*N*_e_) between approximately 3,000 and 10,000 years ago. We therefore incorporated a population-size-change event along the *pepo* lineage and constrained its timing to this interval, with the timing estimated by momi2. This event was interpreted as a demographic bottleneck potentially associated with domestication rather than as a direct estimate of the wild–cultivated split. The momi2 analysis was designed to estimate major divergence times within this predefined framework rather than to reconstruct the full admixture history inferred by qpGraph. Model adequacy was assessed by simulating datasets under the fitted demographic model using msprime^26^ (v1.4.2). Pairwise 2D SFS were calculated using the same projection scheme as for the empirical data. Empirical data were subsampled to obtain numbers of segregating sites comparable to those in the simulated datasets, and simulated spectra were scaled to the corresponding observed totals.

The ages of SNPs and SVs were estimated using GEVA^40^ (v1beta), assuming a mutation rate of 6.5 × 10⁻^9^ per site per generation. Because GEVA requires a single effective population size, we used the mean *N*_e_ (51,278) inferred by SMC++ for subsp. *pepo* over the interval from 100 to 100,000 years ago. A genome-wide average recombination rate was estimated from a published *C. pepo* genetic map with a total length of 2,199.098 cM^86^. Dividing the total genetic length by the total physical length of the 20 chromosomes in the C39 reference genome yielded an average recombination rate of 6.09 × 10⁻^8^ per bp per generation. Variant ages were estimated using GEVA’s joint clock model, which integrates mutation- and recombination-based information. To evaluate the accuracy of allele-age estimates under the demographic history representative of *C. pepo*, we additionally applied the same GEVA parameters to variants simulated with msprime under the inferred demographic model and compared the GEVA-estimated ages with their known simulated mutation ages.

### Distribution of fitness effects calculation

Ancestral and derived alleles for each variant were determined using watermelon (*Citrullus lanatus*)^61^ and *Cucurbita argyrosperma*^87^‒the former distantly related and the latter closely related to *Cucurbita pepo*‒as outgroup species. SNP and SV alleles shared among *C. pepo*, *C. lanatus*, and *C. argyrosperma* were considered ancestral. The distribution of fitness effects (DFE) and the proportion of adaptive variants (α) for nSNPs and SVs were estimated using polyDFE^27^ (v2.15) based on the unfolded site frequency spectrum (SFS). The mean α estimates were derived from discretized DFEs inferred through 20 bootstrap replicates.

### Climatic variables collection and redundancy analysis

Climatic variables were obtained from the WorldClim database (https://www.worldclim.org/), which provides high-resolution global climate surfaces derived from long-term weather station records. Nineteen standard bioclimatic variables (BIO1–BIO19), including 11 temperature-related (BIO1–BIO11) and 8 precipitation-related variables (BIO12–BIO19), were downloaded at a spatial resolution of 30 arc-seconds (∼1 km²), representing average climatic conditions for 1970– 2000. Future climate projections were obtained from downscaled CMIP5 climate data available under the RCP8.5 greenhouse gas emission scenario, representing a high-emission trajectory^88^. Bioclimatic variables for two future periods‒mid-century (2040-2060) and late-century (2080-2100)‒were downloaded from the WorldClim database to assess potential environmental variation across the geographic range of *C. pepo*.

Geographic coordinates for each accession were used to extract corresponding climatic values from raster layers using the extract function in the R package raster (v3.3.13) (https://cran.r-project.org/web/packages/raster). To evaluate the relationship between genomic variation and climatic gradients, we performed redundancy analysis (RDA)^34^ using the R package vegan (https://cran.r-project.org/web/packages/vegan/index.html). RDA was conducted separately for SNPs and SVs, and only variants without missing data and with minor allele frequency (MAF) > 0.05 were retained as response variables to minimize the influence of rare alleles. The 19 bioclimatic variables were used as explanatory variables in the RDA models. To quantify the contribution of climatic predictors to genomic variation, we further conducted variance partitioning within the RDA framework to estimate the proportion of genetic variation explained by the full set of climatic variables. Statistical significance of the RDA models was assessed using permutation tests implemented in the vegan package.

### Genome–environment association analysis

Associations between environmental variables and genome-wide variants were identified using Bayenv2^36^ with the 19 bioclimatic variables. To account for population structure, a covariance matrix describing population allele frequency covariance was estimated using 100,000 MCMC iterations. For each SNP or SV, Bayes factors (BFs) were computed across 10,000 iterations to evaluate associations between allele frequencies and environmental variables. To ensure robustness, the analysis was repeated five times, and the median BF across runs was used as the final statistic for each variant. Empirical null BF distributions were generated by permuting environmental variables across populations. Variants with BF values exceeding the top 0.1% of the permutation-based BF distribution for each environmental variable were retained as climate-associated loci.

### Species distribution modelling

Species distribution models (SDMs)^42^ were constructed using the R package dismo (https://cran.r-project.org/web/packages/dismo/index.html) to estimate the potential habitat range of *C. pepo* under current and future climate conditions. For wild populations, occurrence data comprised the sampling localities of 12 accessions in this study and georeferenced records for additional 40 accessions obtained from the US National Plant Germplasm System (NPGS) database (https://npgsweb.ars-grin.gov/gringlobal/) (**Supplementary Table 15**). For cultivated populations, geographic coordinates for 116 accessions in this study were used. The 19 bioclimatic variables were used as environmental predictors, and BIOCLIM models were fitted separately for each population.

Model performance was evaluated using five-fold cross-validation. Occurrence and background records were partitioned into five subsets, with four subsets used for model training and the remaining subset used for testing in each iteration. Predictive performance was assessed using the area under the receiver operating characteristic curve (AUC) and the true skill statistic (TSS), with means and standard deviations calculated across the five folds. To assess the robustness of projected range changes to variation in occurrence sampling, we further performed a jackknife resampling analysis. For each population, models were independently refitted 20 times, each using a randomly selected 80% subset of the occurrence records. Each refitted model was projected under current climate conditions and future climate scenarios for mid-century (2040-2060) and late-century (2080-2100), and the percentage change in suitable habitat area relative to the current distribution was calculated. The mean and standard deviation of projected range changes across the 20 jackknife replicates were used to evaluate the stability of the projections. Final models were projected under current and future climate scenarios for mid-century and late-century periods.

### Estimating genetic offset under future climates

Genetic offset under future climate scenarios was estimated using Gradient Forest modeling^37^. Climate-associated SNPs and SVs identified from genome–environment association analyses were used as response variables, and bioclimatic variables were used as environmental predictors. Gradient Forest modeling was conducted using the R package gradientForest (https://gradientforest.r-forge.r-project.org/). The fitted models were then used to predict genomic turnover under both current and future climatic conditions. Future climate layers representing mid-century (2040–2060) and late-century (2080–2100) conditions were projected onto the Gradient Forest models to obtain predicted genomic compositions. Genetic offset was calculated as the Euclidean distance between predicted genomic compositions under current and future climates. Spatial patterns of genetic offset were visualized using the R package rasterVis (https://cran.r-project.org/web/packages/rasterVis/).

### Genetic load and partial correlation analysis

We used the relative burdens of derived nonsynonymous SNPs (nSNPs) and SVs as proxies for genetic load. For each accession, nSNP and SV loads were calculated as the ratio of the number of derived nonsynonymous and SV alleles, respectively, to the number of derived synonymous alleles^89^. To account for potential effects of demographic history on the relationships among genetic load, climate-associated variation, and genetic offset, nucleotide diversity at fourfold-degenerate sites (π_s_) was used as a proxy for long-term effective population size (*N*_e_), based on the expected relationship π_s_ ≈ 4*N*_e_μ under a standard neutral diploid model^43,90^. Accessions were grouped into genetically defined clusters using k-means clustering of the first ten principal components derived from genome-wide SNP variation. π_s_ was calculated separately for each genetic cluster using fourfold-degenerate sites, and each accession was assigned the π_s_ value estimated for its corresponding cluster. Pearson partial correlations were then calculated between genetic load and the number of climate-associated loci, genetic load and genetic offset, and the number of climate-associated loci and genetic offset, while controlling for π_s._

## Data availability

Raw genome resequencing reads have been deposited in the NCBI BioProject database under the accession number PRJNA1422823. Raw HiFi reads and genome assemblies have been deposited in the NCBI BioProject database under the accession number PRJNA1209910. Genome assemblies and annotations, SNPs, small indels and SVs in VCF format are available at CuGenDBv2 (http://cucurbitgenomics.org/v2/ftp/pan-genome/squash/).

## Author contributions

Z.F., Y.X. and C.W. conceived, designed and supervised the study. X.Z. and H.S. contributed to genome assembly and annotation, pangenome construction, and SV genotyping. X.Z. performed population genetic analyses and genome–environment association analyses. K.B., J.Z., A.A.S., A.F., H.S.P., E.O., A.G., R.G., C.W. and Y.X. contributed to sample collection, DNA extraction, and genome sequencing. X.Z. wrote the manuscript. Z.F. revised the manuscript.

## Acknowledgements

The authors thank Michael Mazourek for providing seeds of the *C. pepo* core collection and for critical reading of this manuscript. This research was supported by grants from USDA National Institute of Food and Agriculture Specialty Crop Research Initiative (2015-51181-24285 and 2020-51181-32139) and Science and Technology Innovation Program of the Chinese Academy of Agricultural Science (CAAS-ASTIP-IVFCAAS).

