## Supplementary Figures for "Graph-based pangenome provides insights into the adaptive evolution of *Cucurbita pepo*"

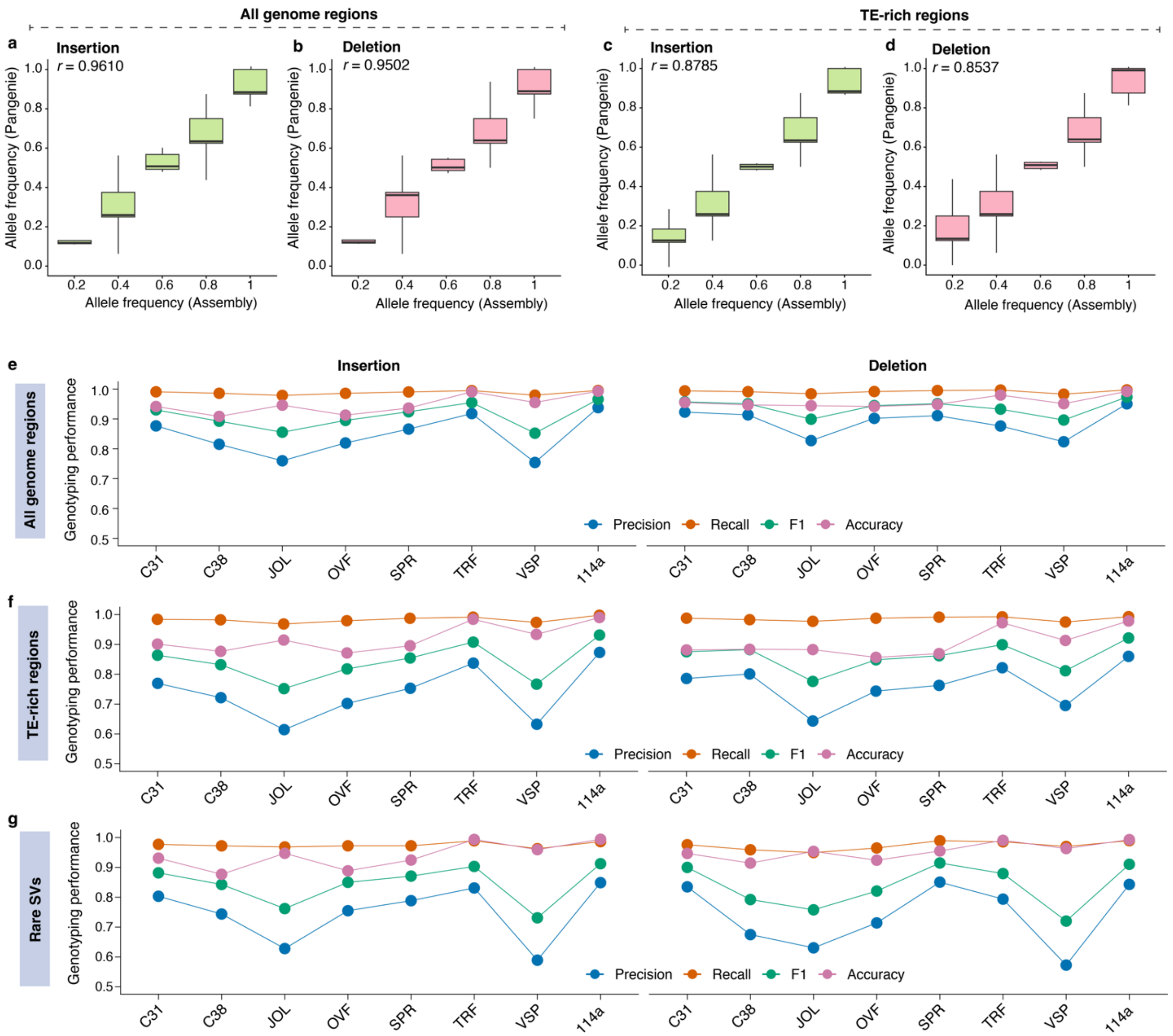


**Supplementary Fig. 1. Validation of graph-based SV genotyping. a–d,** Pearson correlations between allele frequencies estimated using PanGenie and assembly-based genotyping for insertions and deletions across all genomic regions (**a, b**) and TE-rich regions (**c, d**). For each boxplot, the lower and upper bounds indicate the first and third quartiles, respectively, the center line indicates the median, and the whiskers extend to 1.5× the interquartile range. **e–g,** Leave-one-out validation of genotyping precision, recall, F1 score and accuracy across eight accessions for all genome regions (**e**), TE-rich regions (**f**) and rare SVs (**g**), shown separately for insertions and deletions.

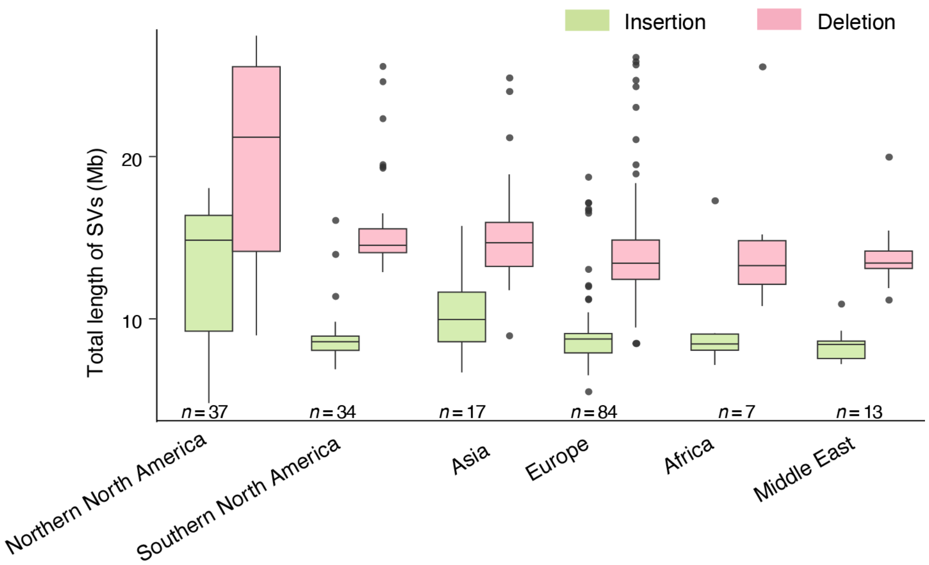


**Supplementary Fig. 2. Total length of SVs across different *C. pepo* populations.** For each boxplot, the lower and upper bounds indicate the first and third quartiles, respectively, the center line indicates the median, and the whiskers extend to 1.5× the interquartile range.

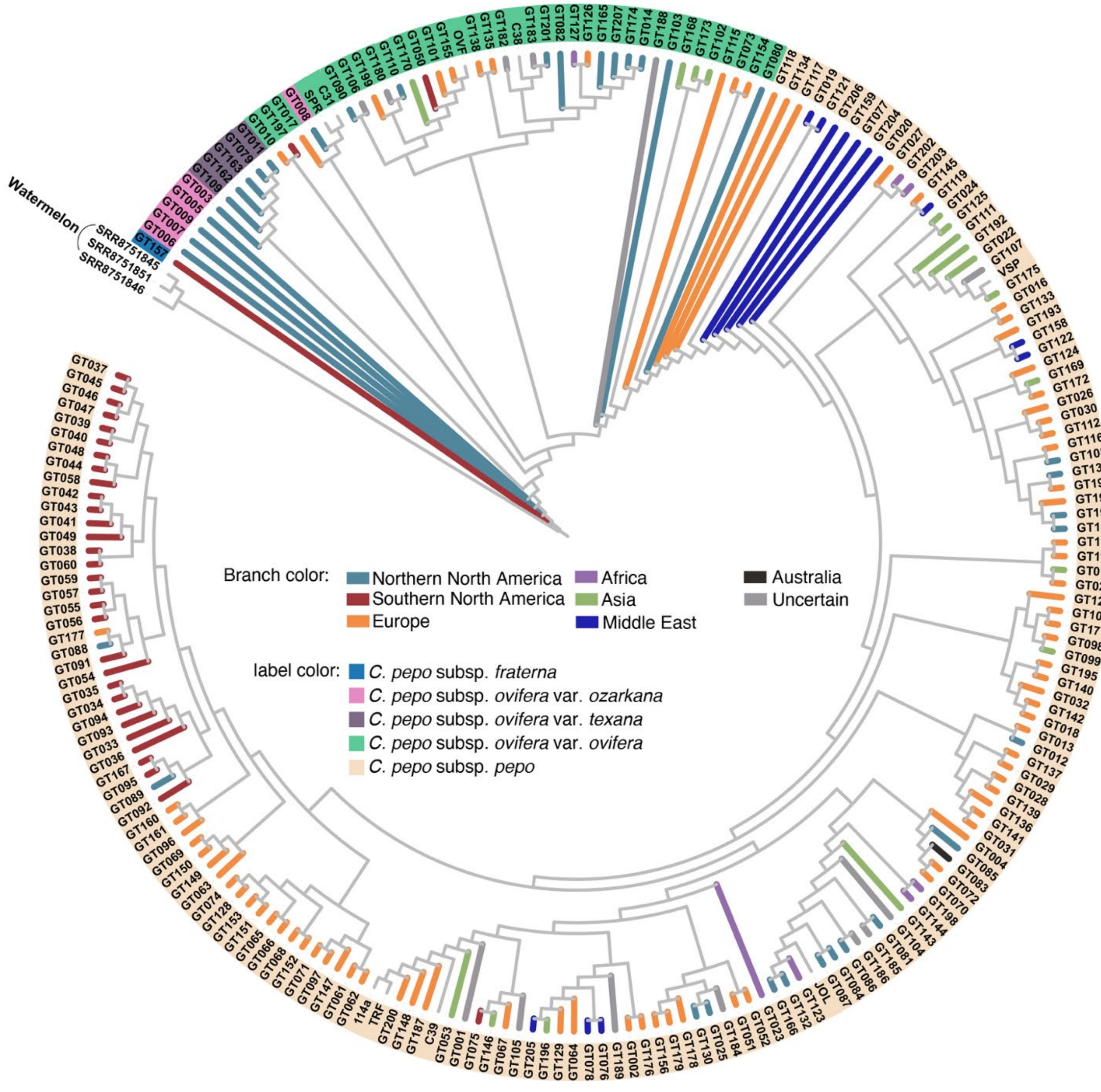


**Supplementary Fig. 3. Phylogenetic relationships among *C. pepo* accessions based on SNPs.**

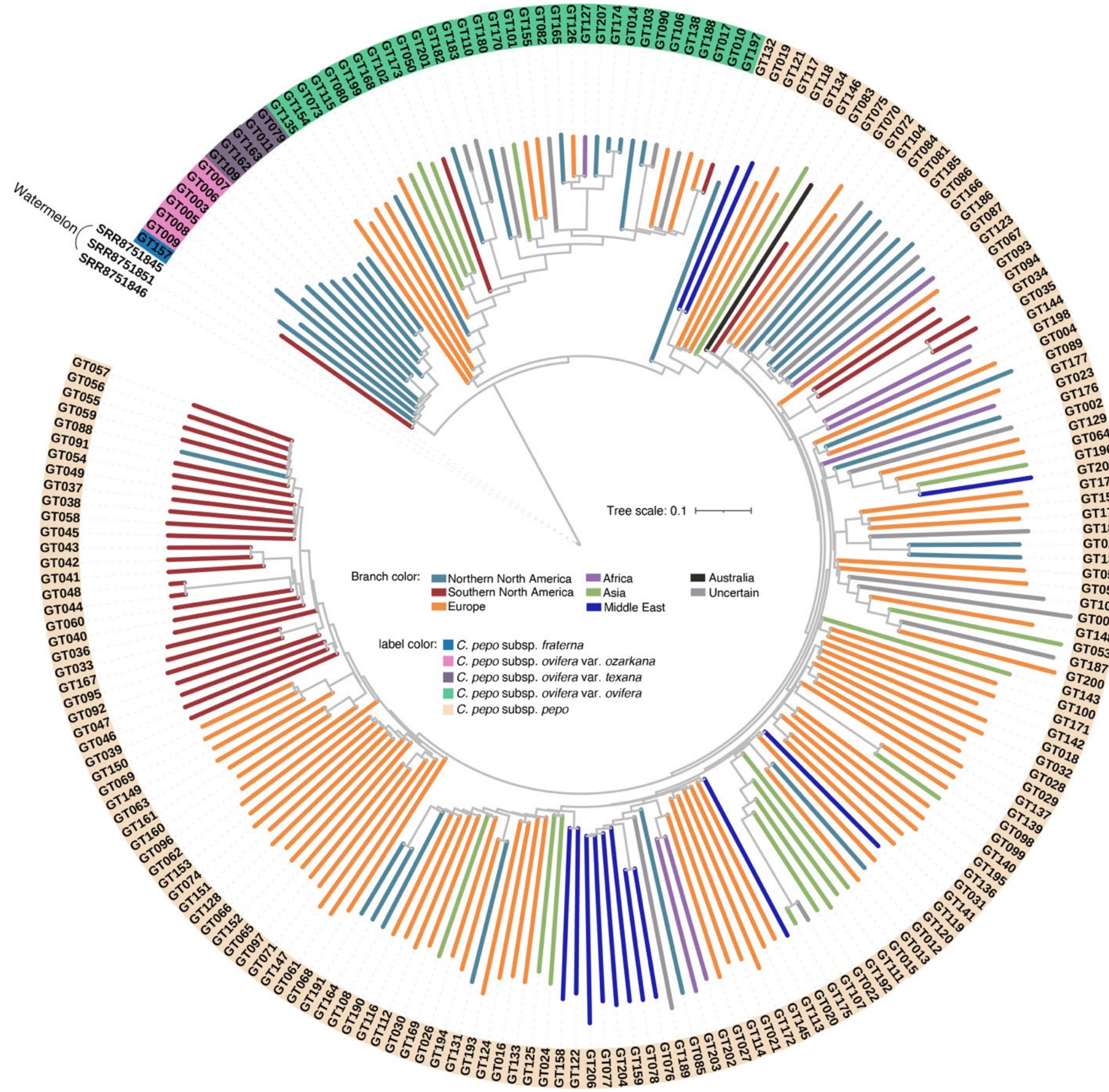


**Supplementary Fig. 4. Phylogenetic relationships among *C. pepo* accessions based on SVs.**

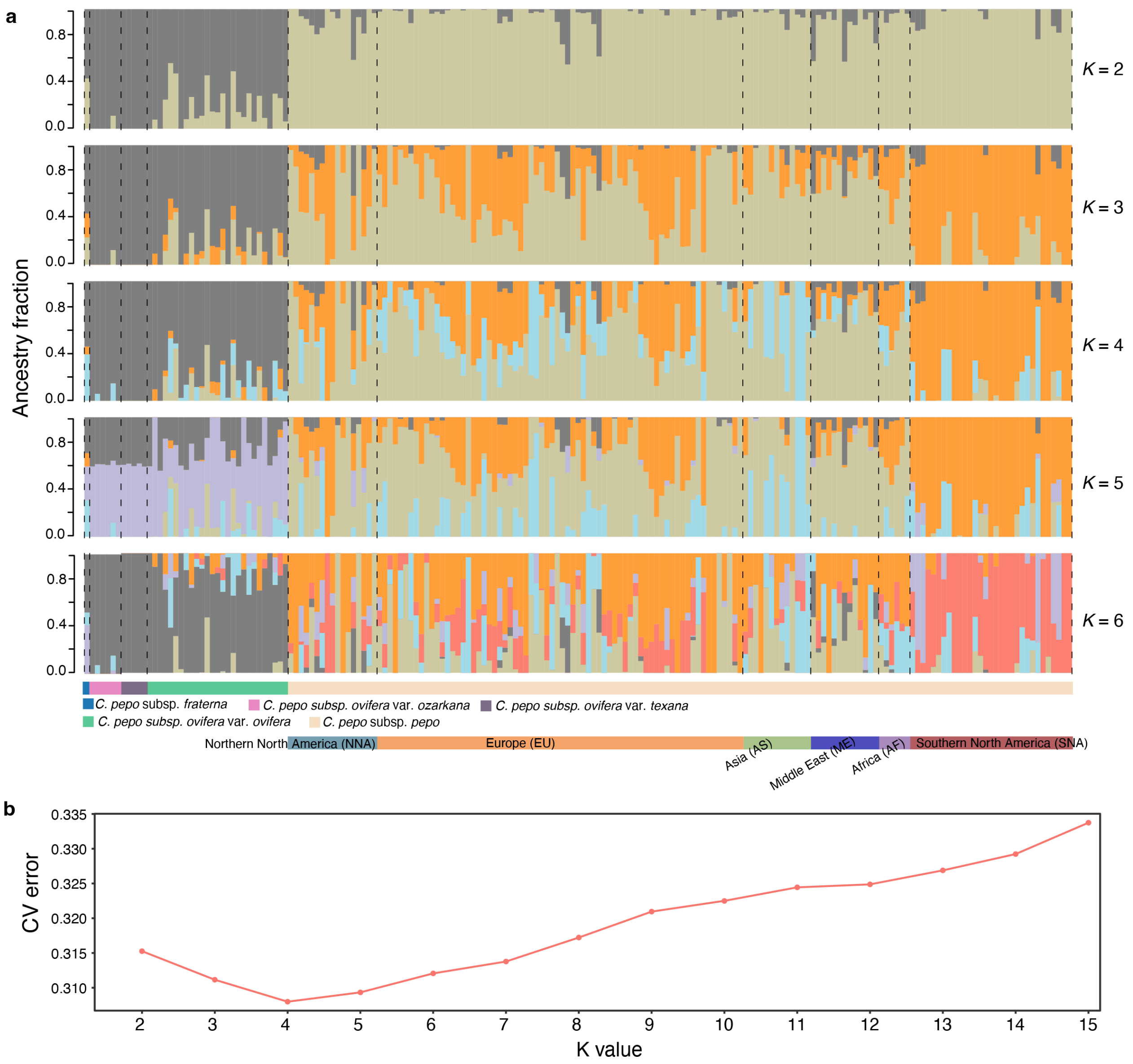


**Supplementary Fig. 5. Population structure of *C. pepo* inferred from SNPs. a,** Bar plots showing ancestry proportions of 206 accessions for *K* values ranging from 2 to 6. Each vertical bar represents one accession, and colors indicate inferred ancestral components. Samples are grouped and ordered by subspecies and geographic origin. **b,** Cross-validation (CV) error estimates from ADMIXTURE runs for *K* = 2–15.

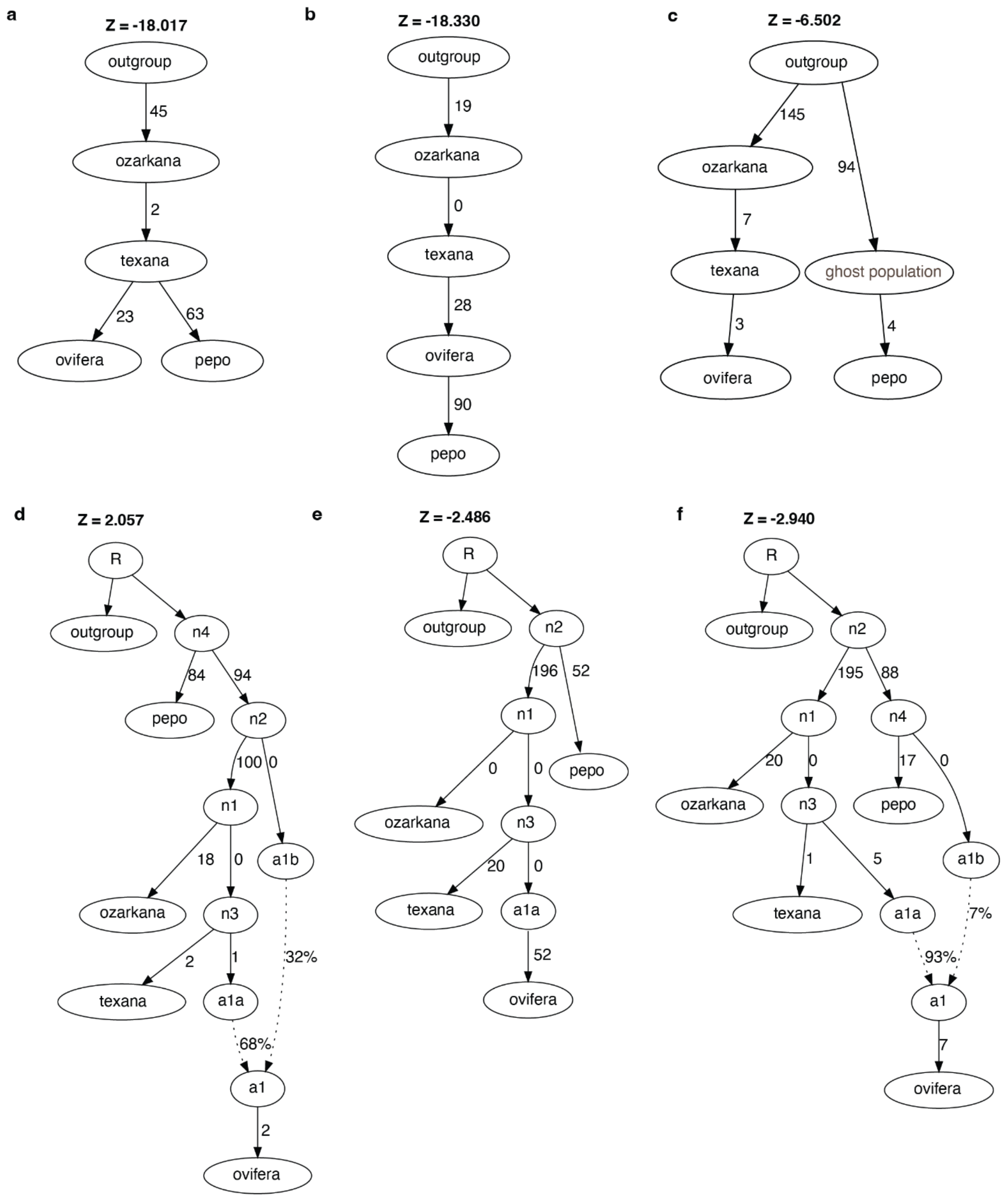


**Supplementary Fig. 6. Admixture graph models. a–c**, Alternative tree models representing different hypotheses of relationships among *C. pepo* groups. **d–f**, Three best-fitting admixture graph models ($|Z| < 3$). Each node represents a population, and arrows indicate ancestral relationships. Numbers on branches denote drift parameters, and dashed arrows indicate admixture events with estimated proportions. Z-scores are shown for each model to indicate goodness of fit. All 92 candidate models and their associated Z-scores are available on GitHub (https://github.com/xuebozhao16/CucurbitGenomics/tree/main/Admixturegraph).

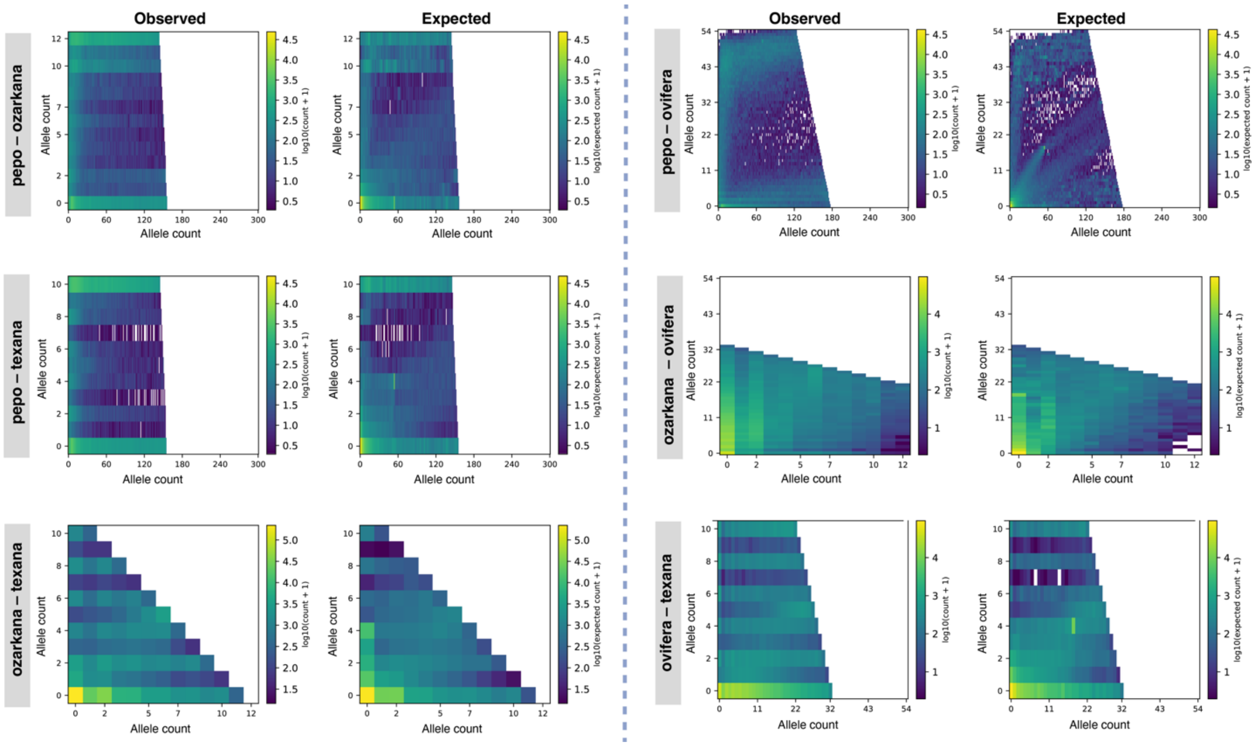


**Supplementary Fig. 7. Comparison of observed and simulated pairwise two-dimensional site-frequency spectra (2D SFS).** Observed 2D SFS (left) and expected 2D SFS simulated using msprime under the inferred demographic model (right) are shown for six population pairs: *pepo–ozarkana*, *pepo–ovifera*, *pepo–texana*, *ozarkana–ovifera*, *ozarkana–texana* and *ovifera–texana*. Colors in the observed and expected spectra represent $\log_{10}$(count+1).

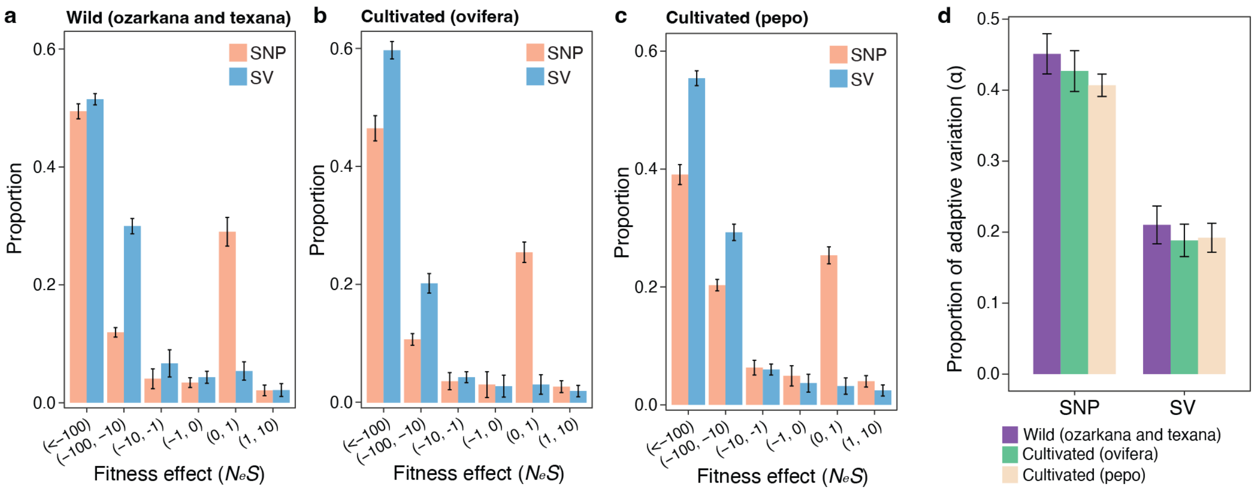


**Supplementary Fig. 8. Distribution of fitness effects and adaptive variation inferred from SNPs and SVs in *C. pepo*. a–c**, Distribution of fitness effects ($N_eS$) for SNPs and SVs in wild (**a**), cultivated *ovifera* (**b**) and cultivated *pepo* (**c**) populations. **d**, Proportion of adaptive variation (α) inferred from SNPs and SVs in wild and two cultivated groups. Values were derived from 20 bootstrap replicates; bars represent mean ± SD.

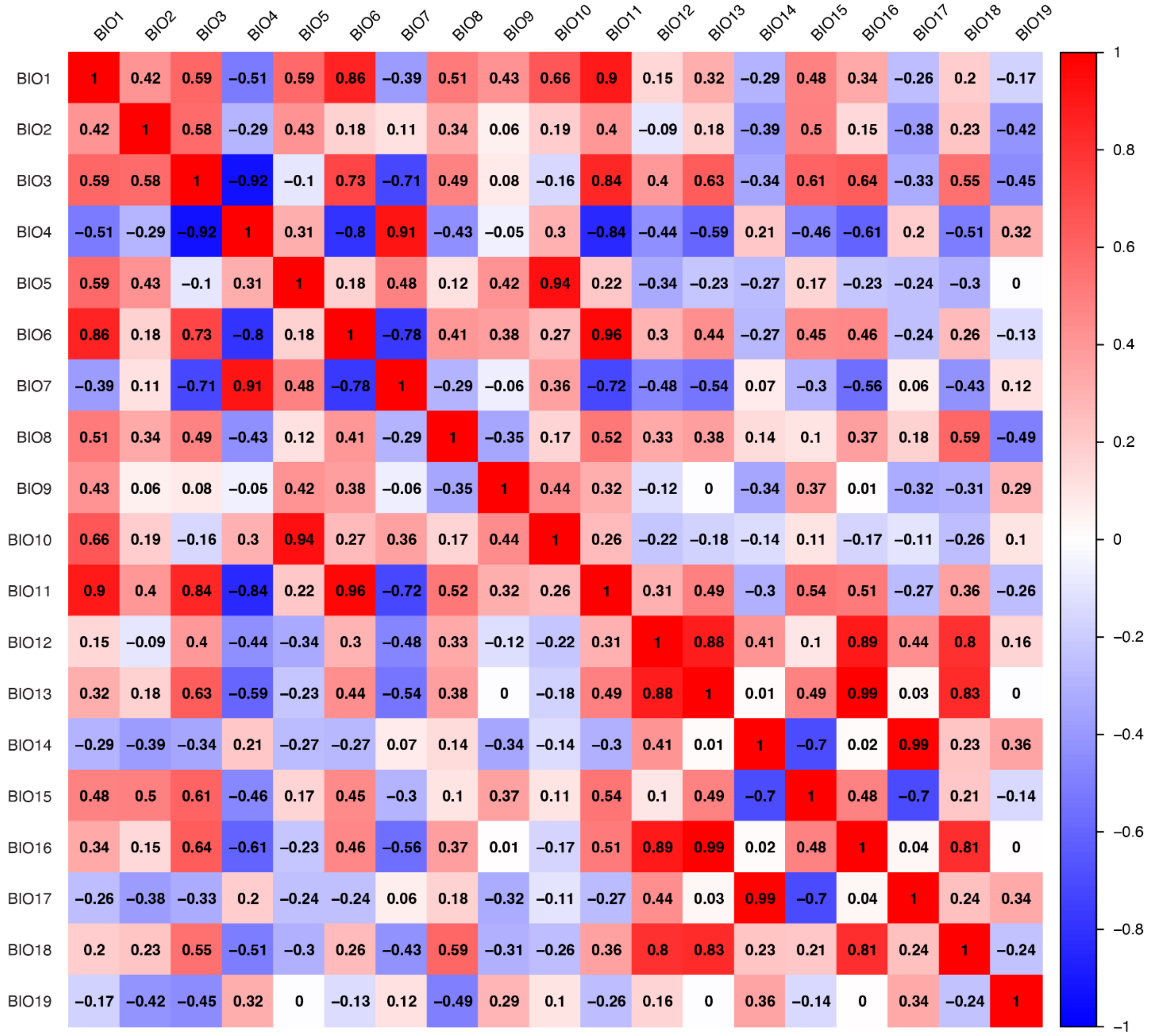


**Supplementary Fig. 9. Correlation matrix of 19 bioclimatic variables.**

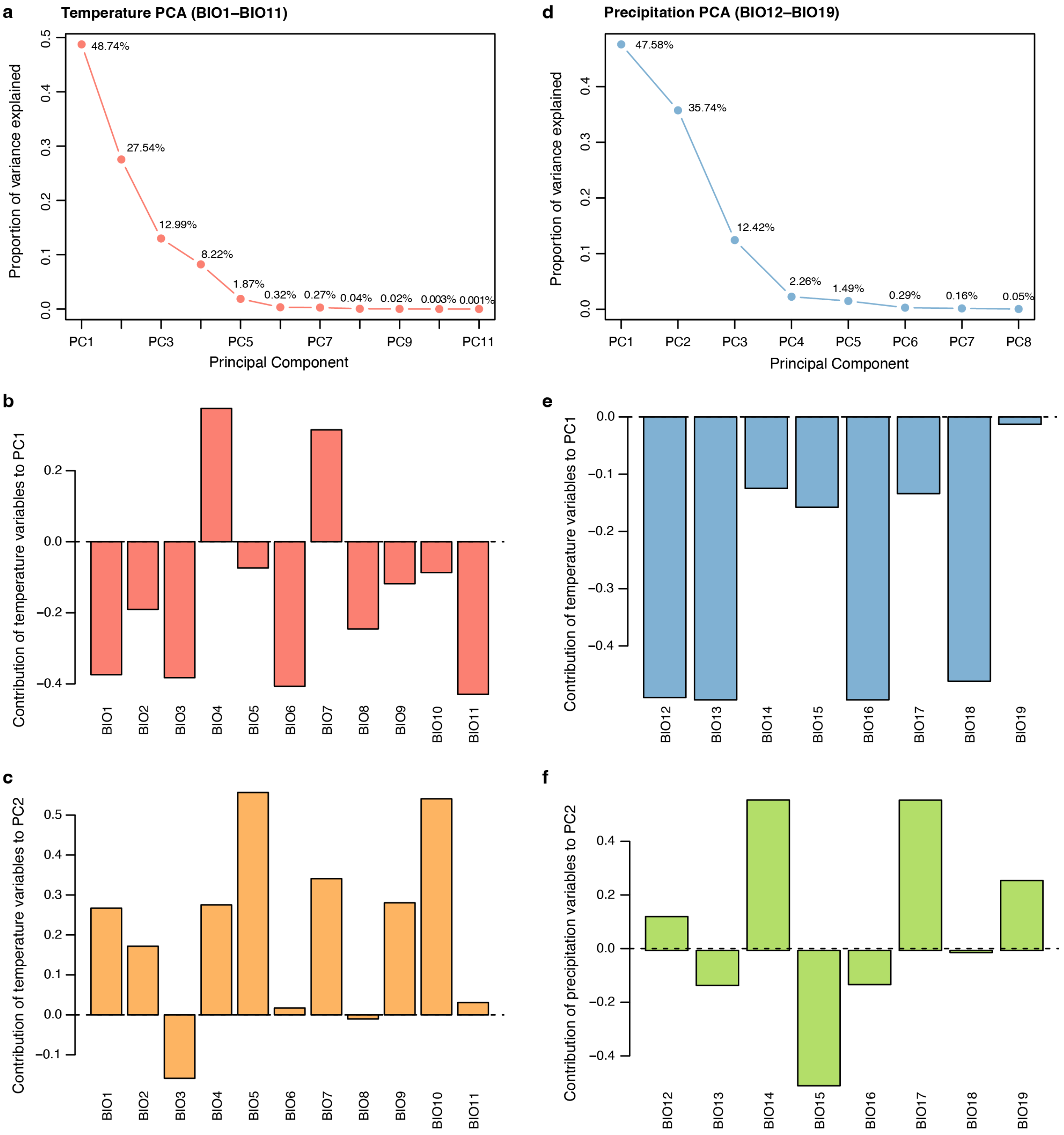


**Supplementary Fig. 10. Principal component analysis (PCA) of temperature- and precipitation-related variables. a,** Percentages of variance explained by principal components derived from PCA of temperature-related bioclimatic variables (BIO1–BIO11). **b,c,** Contribution of individual temperature variables to the first (**b**) and second (**c**) principal components. **d,** Percentages of variance explained by principal components derived from PCA of precipitation-related bioclimatic variables (BIO12–BIO19). **e,f,** Contribution of individual precipitation variables to the first (**e**) and second (**f**) principal components.

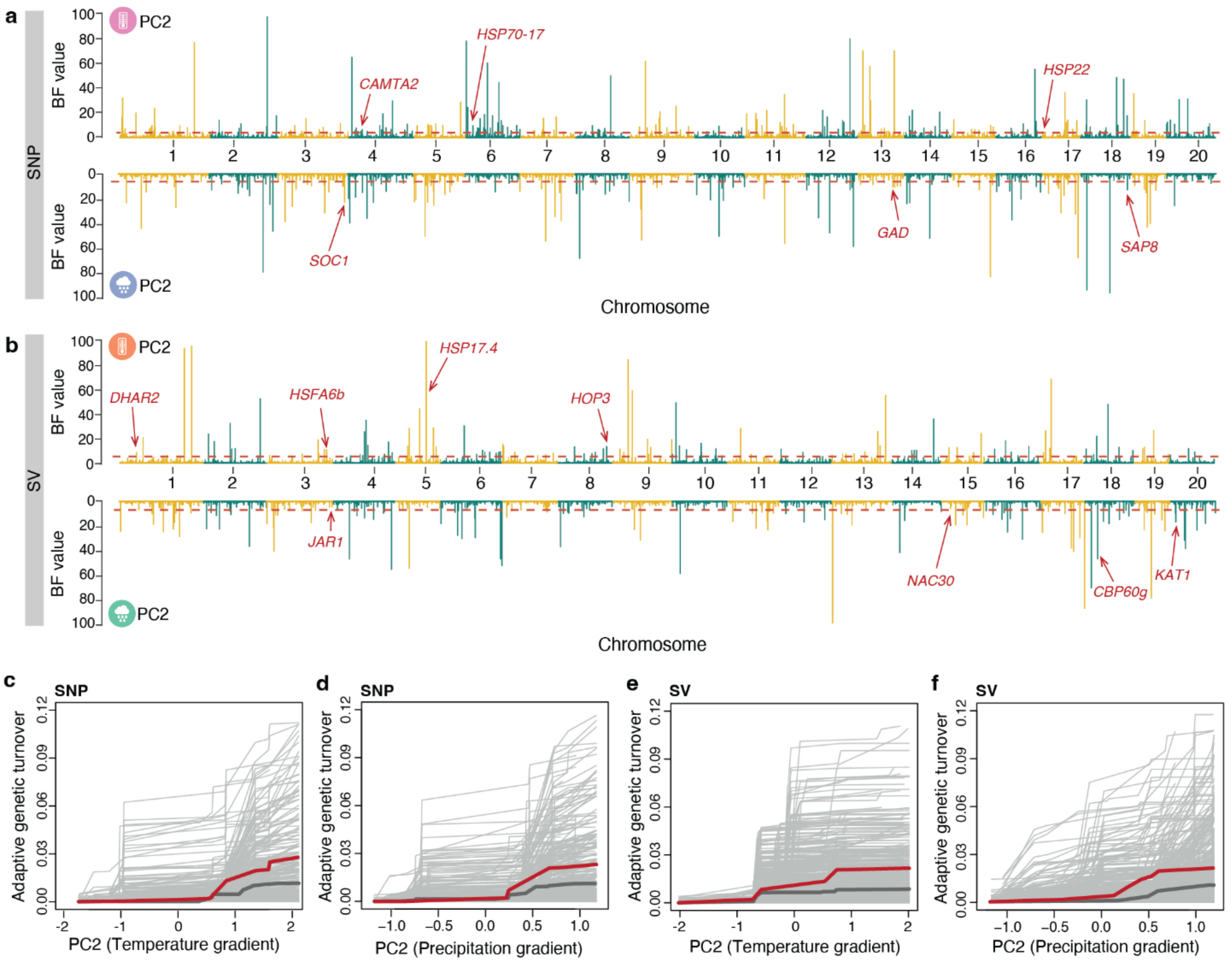


**Supplementary Fig. 11. Climate-associated genetic variation along PC2 of climatic gradients. a,b,** Genome-wide distribution of climate-associated SNPs (**a**) and SVs (**b**) along PC2 of temperature (top panels) and precipitation (bottom panels) gradients. The dashed red line indicates the significance threshold, and representative candidate genes near the top signals are shown. Details of the representative candidate genes are provided in **Supplementary Table 12**. **c–f,** Turnover of adaptive SNPs (**c**,**d**) and SVs (**e**,**f**) along PC2 of temperature and precipitation gradients. Thin light-grey lines represent turnover trajectories of individual loci, thick dark-grey lines indicate the mean cumulative turnover of random loci, and colored thick lines denote the mean turnover of climate-associated adaptive loci.

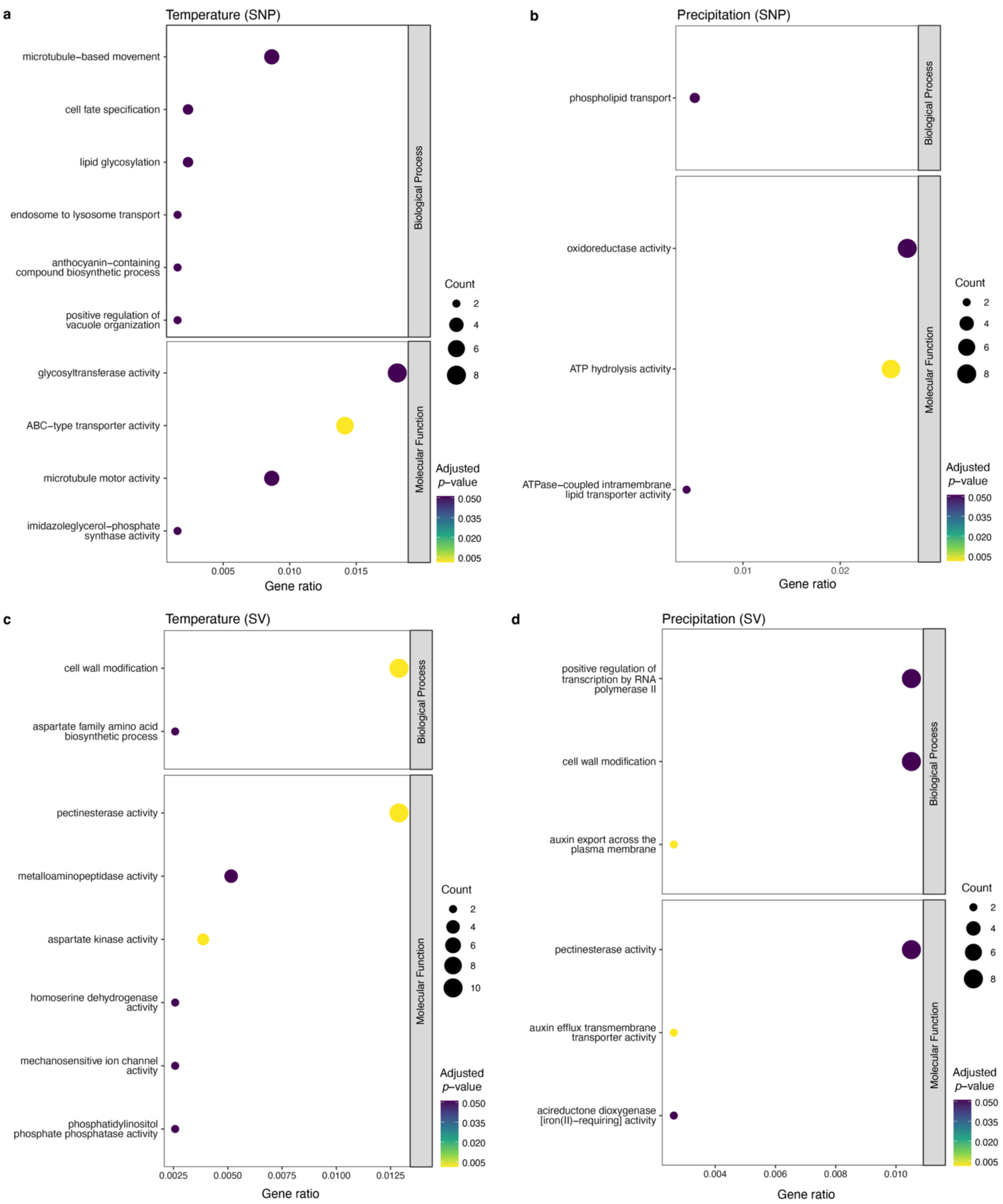


**Supplementary Fig. 12. Functional enrichment of climate-associated candidate genes across variant types (SNPs and SVs) and climatic dimensions (temperature and precipitation).**

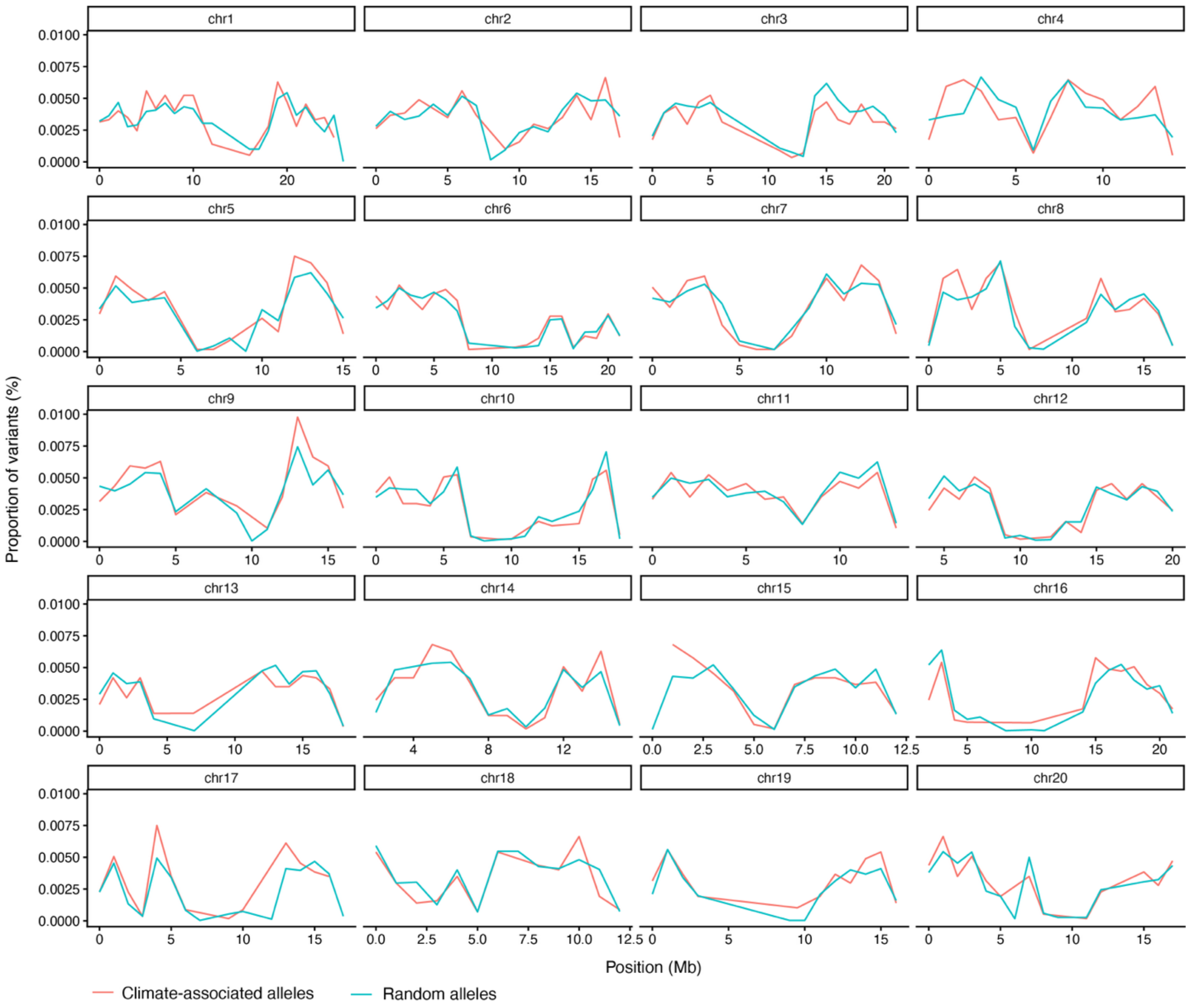


**Supplementary Fig. 13. Genomic distributions of climate-associated and random variants across the *C. pepo* genome.** Climate-associated and random variants were grouped into 1-Mb non-overlapping genomic windows. For each variant class, the proportion of variants located within each window was calculated and plotted along each chromosome.

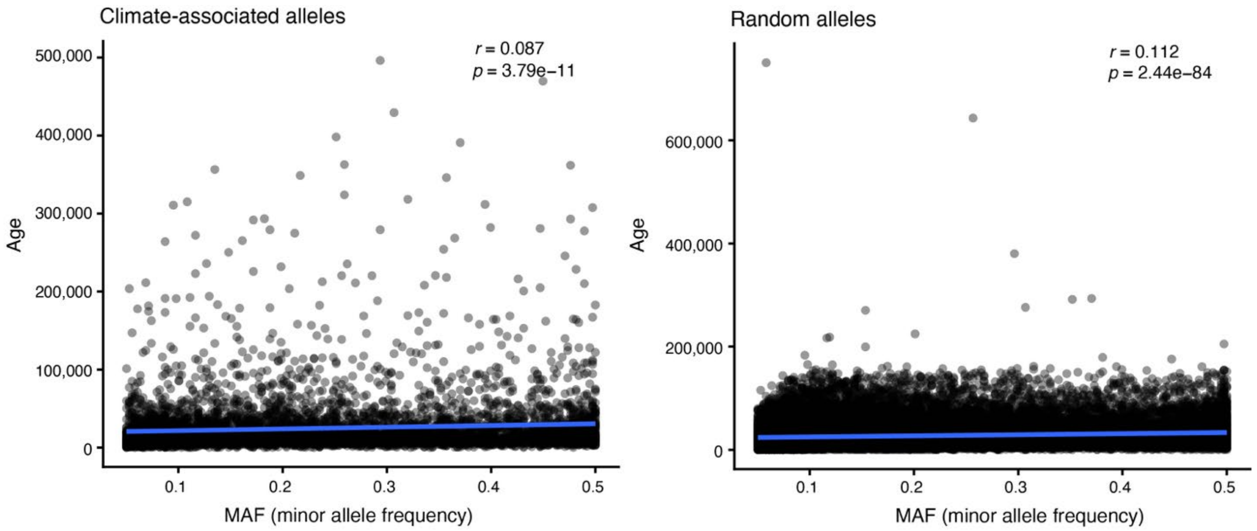


**Supplementary Fig. 14. Relationship between minor allele frequency and estimated allele age.** Minor allele frequency (MAF) is plotted against estimated allele age for climate-associated SNPs (left) and randomly sampled background SNPs (right). Each point represents an individual SNP, and blue lines indicate fitted linear regressions. Pearson correlation coefficients (*r*) and corresponding *p* values are shown in each panel.

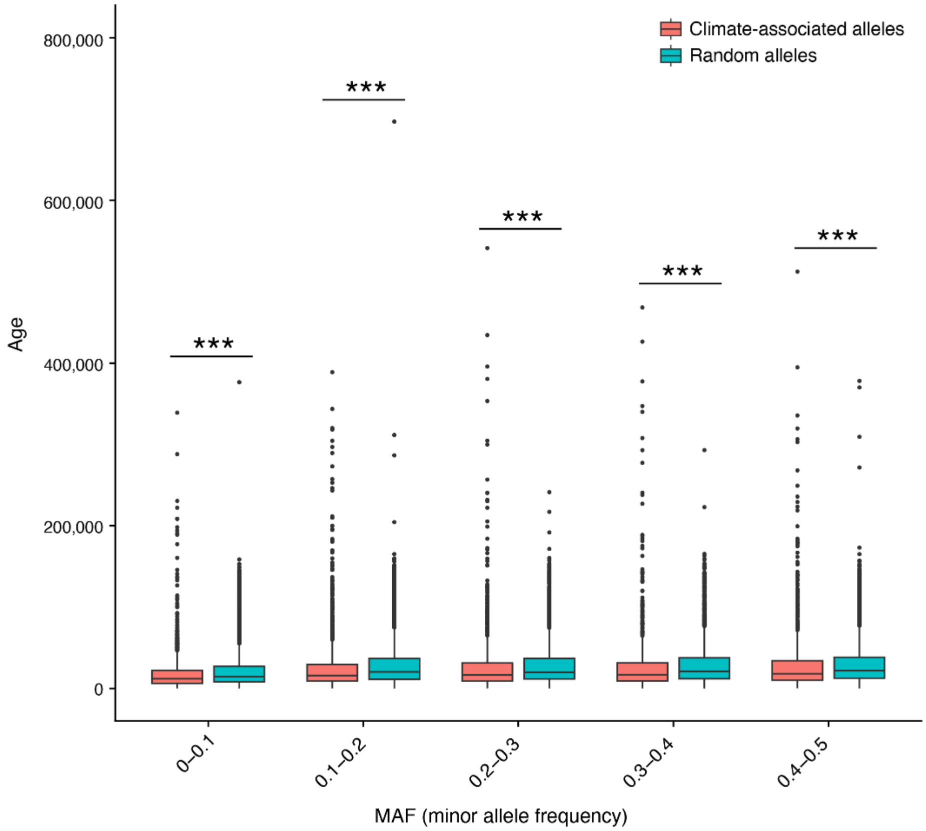


**Supplementary Fig. 15. Allele ages of climate-associated and random SNPs across minor allele frequency classes.** Estimated allele ages of climate-associated and randomly sampled background SNPs are compared across five minor allele frequency (MAF) classes. For each boxplot, the lower and upper bounds indicate the first and third quartiles, respectively, the center line indicates the median, and the whiskers extend to 1.5× the interquartile range. Asterisks indicate significant differences in allele age between climate-associated and random SNPs within each MAF class ($p < 0.001$; Wilcoxon rank-sum test).

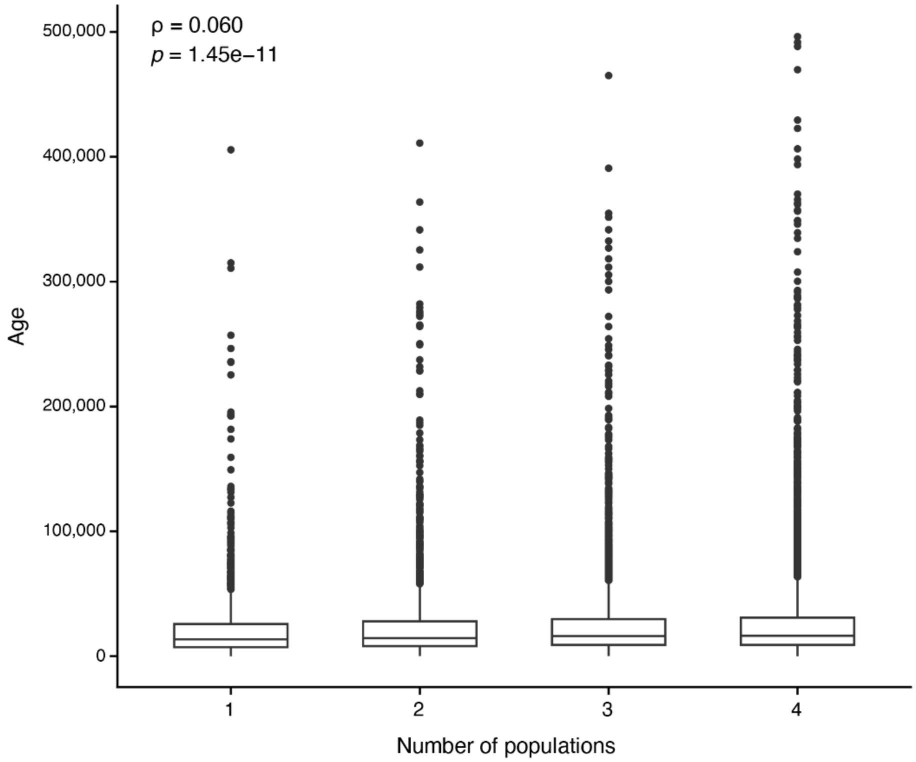


**Supplementary Fig. 16. Relationship between allele age and population sharing of climate-associated alleles.** Climate-associated SNPs were grouped according to the number of major populations in which the allele was present. Boxplots show the distributions of estimated allele ages for SNPs shared across one to four populations. The Spearman rank correlation coefficient (ρ) between allele age and the number of populations in which the alleles were present, together with the corresponding *P* value, is shown. For each boxplot, the lower and upper bounds indicate the first and third quartiles, respectively, the center line indicates the median, and the whiskers extend to 1.5× the interquartile range.

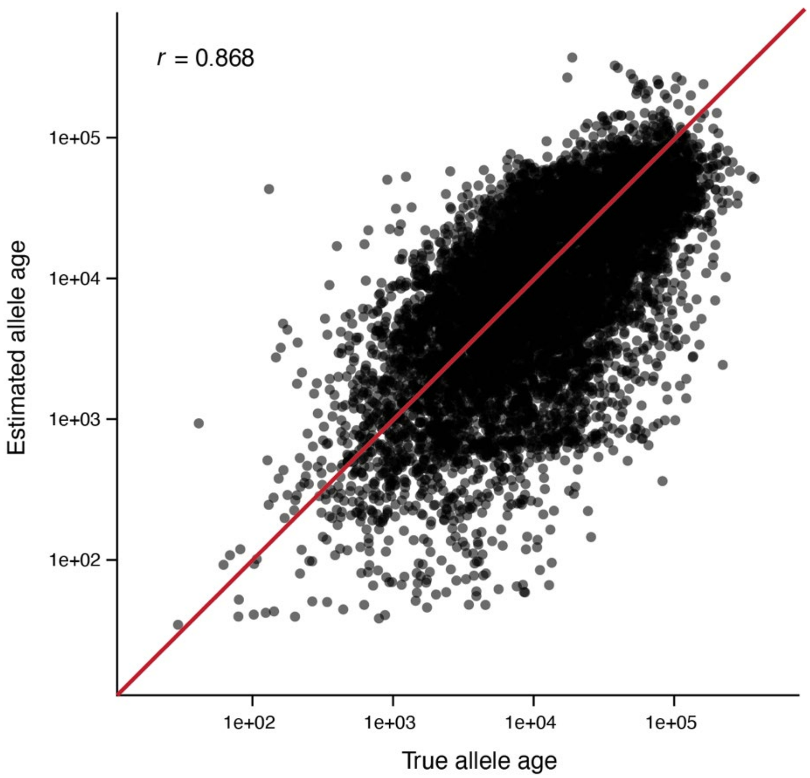


**Supplementary Fig. 17. Validation of allele-age estimates using simulated data.** GEVA-estimated allele ages were compared with the true ages of variants simulated under the inferred demographic model. Each point represents an individual simulated variant, with both axes shown on a logarithmic scale. The red line indicates the 1:1 relationship between estimated and true allele ages. The Pearson correlation coefficient (*r*) between estimated and true allele ages is shown.

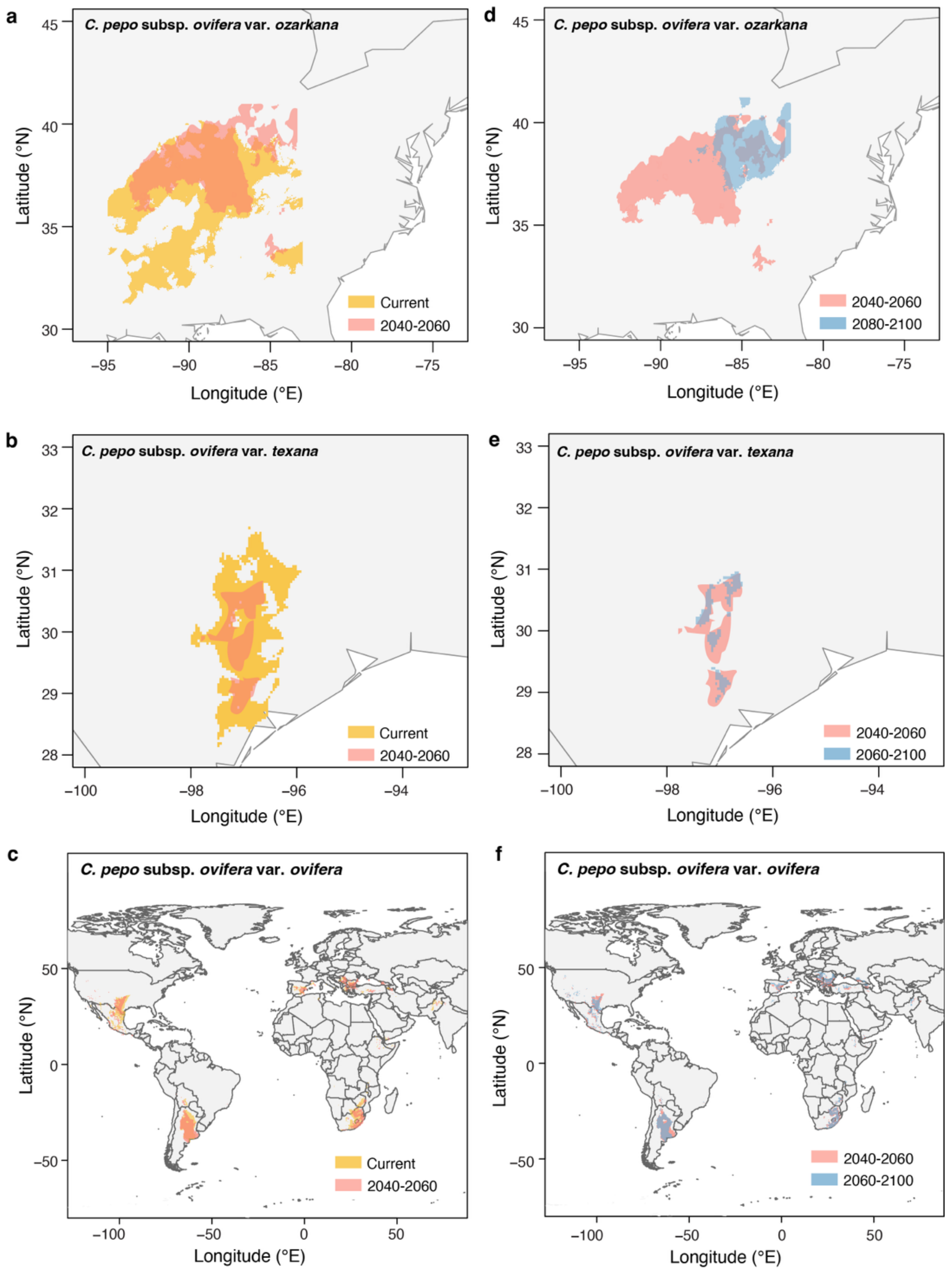


**Supplementary Fig. 18. Projected climatic suitability of *C. pepo* subsp. *ovifera* taxa under future climate scenarios. a–c,** Predicted climatically suitable areas for *C. pepo* subsp. *ovifera* var. *ozarkana* (**a**), var. *texana* (**b**) and var. *ovifera* (**c**) under current and mid-century (2040–2060) climate conditions. **d–f,** Comparison of projected suitable habitats for *C. pepo* subsp. *ovifera* var. *ozarkana* (**d**), var. *texana* (**e**) and var. *ovifera* (**f**) between mid-century (2040–2060) and late-century (2080–2100) climate scenarios. Maps were created using the R package rworldmap (https://cran.r-project.org/web/packages/rworldmap/index.html).

**a**

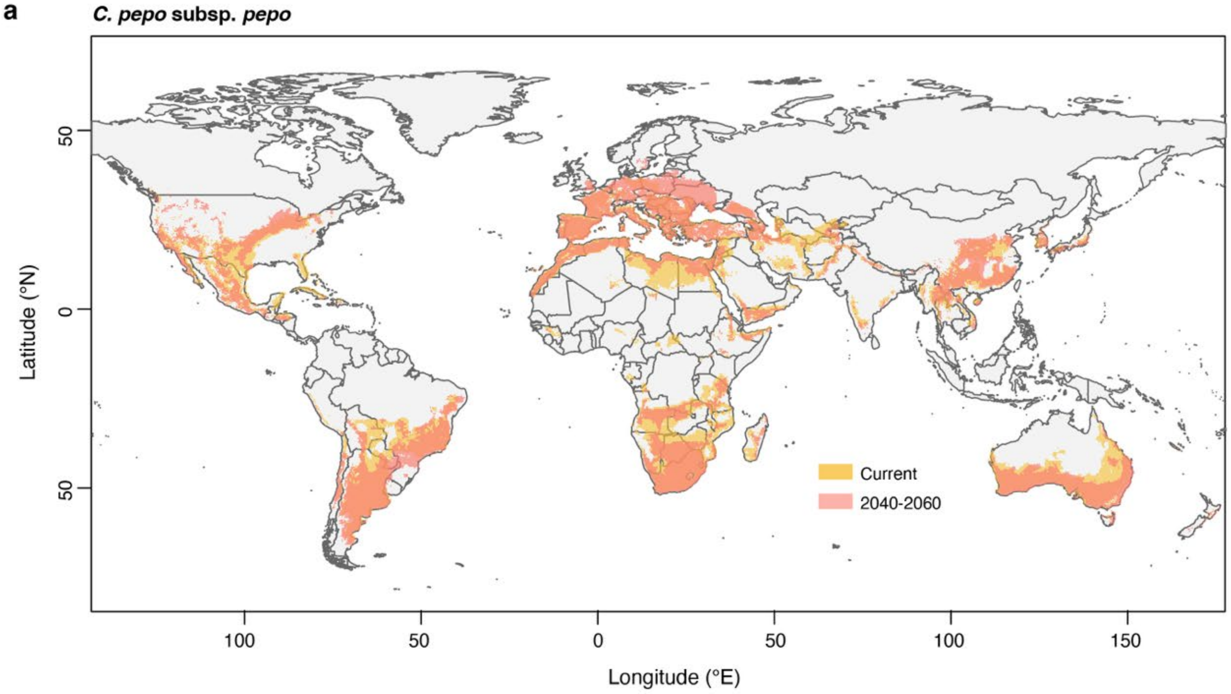


**b**

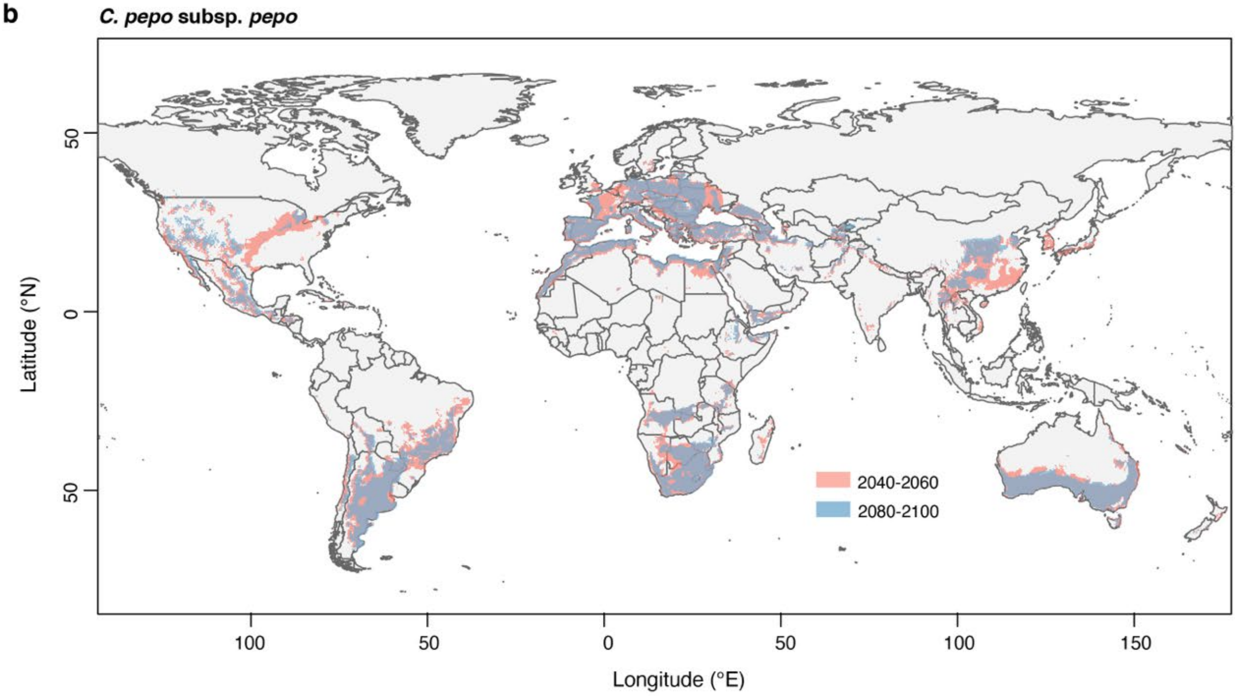


**Supplementary Fig. 19. Projected climatic suitability of *C. pepo* subsp. *pepo* under future climate scenarios: mid-century (2040–2060) and late-century (2080–2100).**

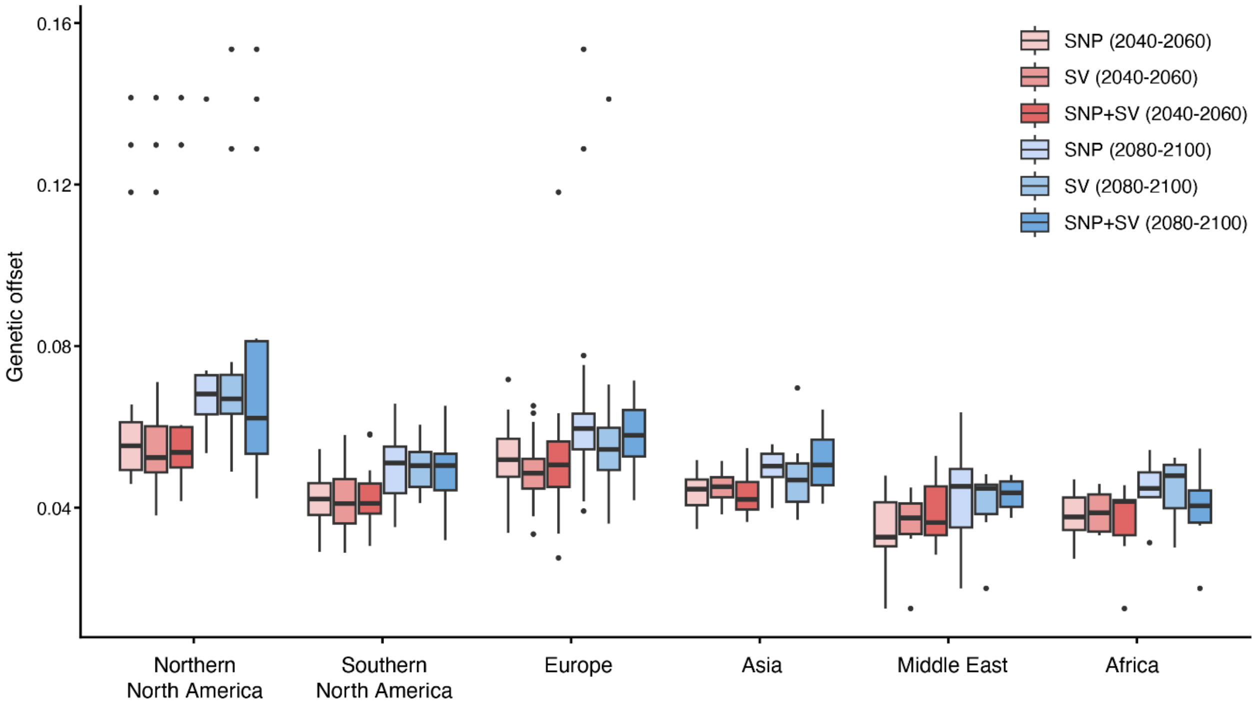


**Supplementary Fig. 20. Regional genetic offset estimated from climate-associated SNPs, SVs, and their combination under mid-century (2040–2060) and late-century (2080–2100) climate scenarios.** For each boxplot, the lower and upper bounds indicate the first and third quartiles, respectively, the center line indicates the median, and the whiskers extend to 1.5× the interquartile range.

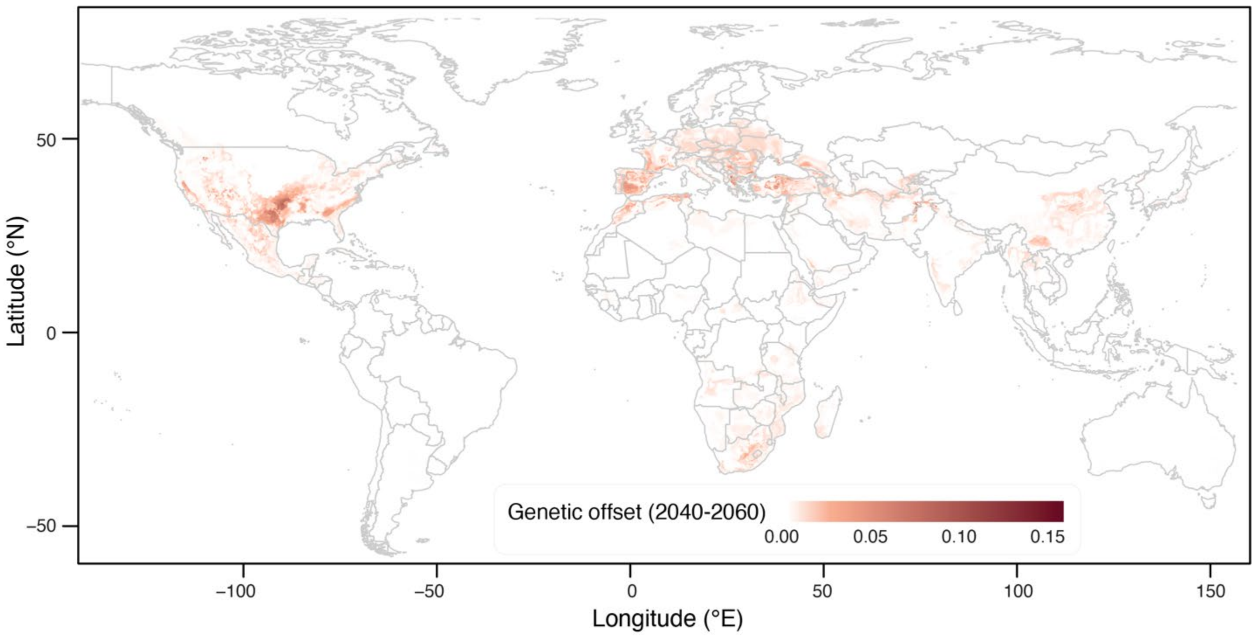


**Supplementary Fig. 21. Spatial distribution of genetic offset projected for mid-century (2040–2060) for the entire *C. pepo* population.**

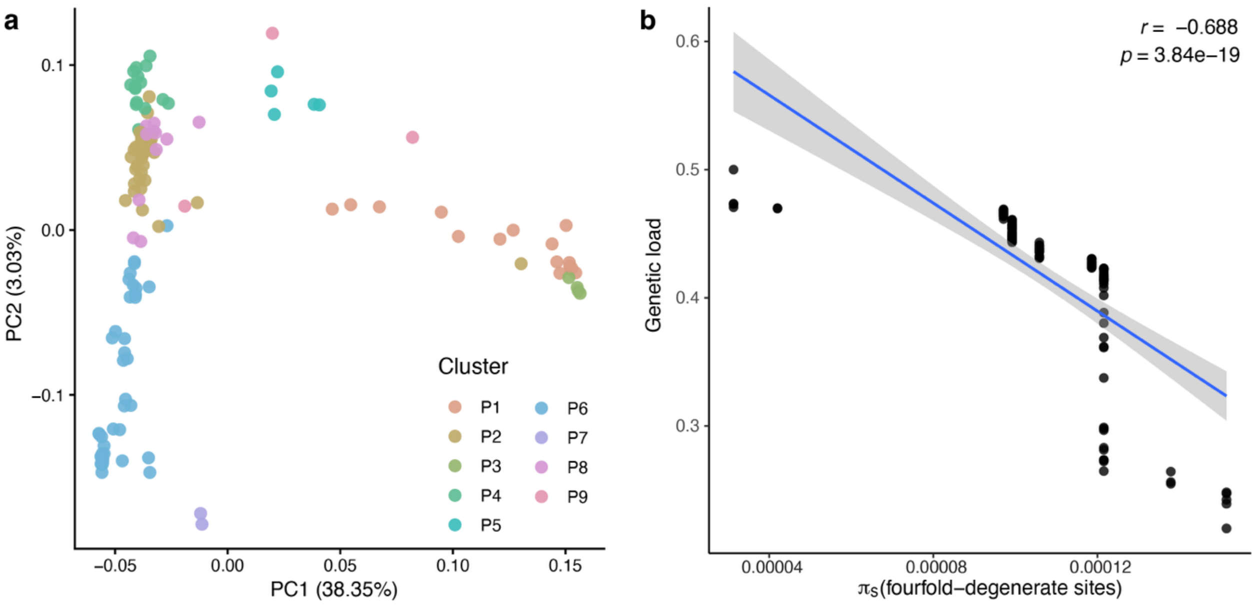


**Supplementary Fig. 22. Genetic structure and relationship between genetic load and nucleotide diversity at fourfold-degenerate sites ($\pi_s$). a,** Principal component analysis of genome-wide SNP variation. Accessions are colored according to 9 genetic clusters identified by k-means clustering. **b,** Relationship between genetic load and $\pi_s$, used as a proxy for long-term effective population size. Each point represents an accession, with the accessions assigned the $\pi_s$ value estimated for its corresponding genetic cluster. The blue line indicates the linear regression, and the shaded area represents the 95% confidence interval. Pearson's correlation coefficient (*r*) and the corresponding *P* value are shown.
